# Speed Vascular Patterns in the Spatial Navigation System

**DOI:** 10.1101/2025.07.15.663536

**Authors:** Felipe Cybis Pereira, Sebastian H. Castedo, Samuel Le Meur-Diebolt, Nathalie Ialy-Radio, Soumee Bhattacharya, Jeremy Ferrier, Bruno Félix Osmanski, Simona Cocco, Remi Monasson, Sophie Pezet, Mickaël Tanter

## Abstract

The hippocampal formation is central to spatial navigation, hosting neurons that encode position, direction, and speed. Yet, the brain-wide vascular dynamics supporting these processes remain poorly understood, especially during naturalistic behaviors. Here, we adapted functional ultrasound (fUS) imaging to examine how cerebral blood volume (CBV) changes relate to behavioral parameters in freely moving rats. High-resolution imaging of hippocampal-parahippocampal regions during open-field exploration reveals strong correlations between CBV dynamics and animal speed, with distinct regional activation patterns and temporal delays. Lagged general linear modeling uncovers information flow from the thalamus to parahippocampal regions, including the medial entorhinal cortex, and to hippocampal subfields (dentate gyrus, CA1–CA3), consistent with a hierarchical processing framework. The analysis also links CBV with angular head speed and the dorsal thalamus.

Decoding analyses show that CBV signals not only encode speed precisely but also capture spatial features like proximity to walls and corners, even when univariate analyses do not. This decoding remains robust across animals, underscoring the universality of speed encoding in vascular dynamics. We also identify slow CBV oscillations in the hippocampus aligned with minute-scale speed fluctuations, suggesting a neurovascular signature of exploratory behavior.

These findings reveal a hemodynamic signature of speed representation in the navigation system, arising from energy demands in a continuous attractor network model for path integration, where population activity and synaptic currents increase quadratically with animal speed as both peak firing rates and neuronal recruitment scale linearly with animal speed. Moreover, they highlight functional ultrasound imaging as a powerful approach for probing the hemodynamic basis of navigation.

## INTRODUCTION

Since the discovery of the brain spatial navigational system, research has revealed the existence of multiple types of spatially-tuned neurons across several species in the hippocampal-entorhinal circuit^1–5^. Neurons called “place cells” located in the hippocampus fire when the animal is in a specific position within an environment, a place field^6,7^, while head direction cells, first discovered in the postsubiculum, fire when the animal is heading to a specific direction, as a brain compass^8–10^. Boundary cells located within the subiculum and entorhinal cortex fire at limiting distance from the environment boundaries^11–13^, while grid cells in the entorhinal cortex fire in multiple place fields, with their position creating a striking regular triangular pattern tessellating the environment^14,15^.

The regularity in space of grid cells firing was rapidly thought as the neuronal substrate providing internal metric for space, a much needed information for path integration models^1,5^. Path integration is the process of calculating one’s current position by integrating self-motion cues, like speed and direction, over time. While place, grid, and head-direction cells provide positional and directional information, their derivative in time, locomotion speed and angular head speed, are critical factors in dynamically updating self-representation in space^2,16,17^. Recently, speed cells have been identified in the medial entorhinal cortex (MEC)^18^, the pre- and parasubiculum (PrS and PaS)^19^ and in the dorsal CA1 in the hippocampus^20^. In a relatively small proportion compared to the other cells of the spatial navigation system, speed cells exhibit firing rates proportional to the animal’s running speed adding a dynamic element to the core spatial representation.

At the mesoscopic scale, individual and local theta rhythm in the hippocampal formation play a role on firing rate timings across spatially-tuned cells^21–24^, and in certain cases, theta rhythm frequency is modulated by running speed, in a weaker but additional layer of speed encoding^25–31^.

The complex relationship between individual neurons, like speed cells, and mesoscale signals like hippocampal theta rhythm and how information flows across the spatial navigation system securing reliable path integration is not yet fully understood^2,3^. Beyond electrophysiological signals, the link between individual neurons involved in spatial navigation and mesoscale cerebral blood flow related to local metabolic demand is also not understood.

In current computational models, path integration relies on the concept of continuous attractor network (CAN)^32^. In CANs, a collective state of neural activity, called bump, encodes the animal’s position, and is updated from sensory e.g. proprioceptive cues during motion. While indirect evidence for the existence of CANs in various brain areas has accumulated over the years^33^, a one-dimensional CAN, coding for head direction and topologically shaped as a ring, was found in Drosophila^34^. An interesting yet, to our knowledge, understudied question concerns the cost associated with updating internal representations. There are already indications that the interaction between neuronal energy and activity generates interesting phenomenology: developing mice have been shown to possess robust head direction systems; however, their self-motion inputs exhibit reduced signaling activity which could be attributed to the high energetic demands of brain development^35^.

Brain-wide imaging techniques that enable observing spatial navigation activity patterns across multiple regions simultaneously are required to offer new insights into the communication and coordination during complex behavior such as spatial navigation.

Functional ultrasound (fUS) imaging^36^ is particularly well-suited for investigating brain-wide activity^37–39^ in freely moving rodents^40–45^. fUS imaging signal is proportional to the local cerebral blood volume (CBV) and often used as a proxy to brain activity through neurovascular coupling^46–48^. In a previous study, where our team performed fUS imaging in rats running in a linear track for water rewards in both ends^42^, results showed strong hemodynamic modulation during each run (fast timescales) as well as hemodynamic adaptation across runs (slow timescales) where relative CBV in the hippocampus linearly increased while cortical blood flow steadily decreased. Running speed and theta activity remained constant across runs. Due to their position in the rat’s brain, the entorhinal cortex and parahippocampal region were not imaged. Also, position, direction and speed are inherently entangled in a linear track task^20^ restricting some analysis from a spatial navigation stand point.

In the present study, we performed functional ultrasound imaging in awake rats freely exploring an open-field to evaluate reliable mesoscopic-scale vascular patterns across the hippocampal formation during spatial navigation and investigate how well these patterns would encode spatial navigation features. Furthermore, we examined the energetic costs of updating a CAN-based path integration model, aiming to understand the theoretical relationship between CBV and animal speed underlying the correlation patterns observed in our recordings. Finally, this correlation between speed and CBV signal in the spatial navigation regions also explains minute-scale oscillations of CBV detected during open-field exploration.

## RESULTS

### fUS imaging reveals robust correlation of local CBV changes with locomotion and angular head speed during free exploration

To investigate whether the variations of CBV in awake, freely moving rats reflect partial activation patterns of place cells, grid cells, or border cells, CBV changes were quantified during exploratory behavior across 12 different parasagittal imaging planes, each including regions of interest (ROI) known to belong to the navigation system. The animal’s behavior was simultaneously recorded. Analysis through pose estimation software^49–51^ allowed the determination of the following key behavioral parameters: the animal’s *x*-*y* position in the arena (Figure 1B-C), the animal’s velocity in *x* or *y* direction (Figure 1C), speed (Figure 1C), head direction (Figure 1C), its proximity to walls (Supplementary Figure 4), the distance from corners (Supplementary Figure 4), positional and grids cells-like patterns (Supplementary Figure 5).

**Figure 1:**
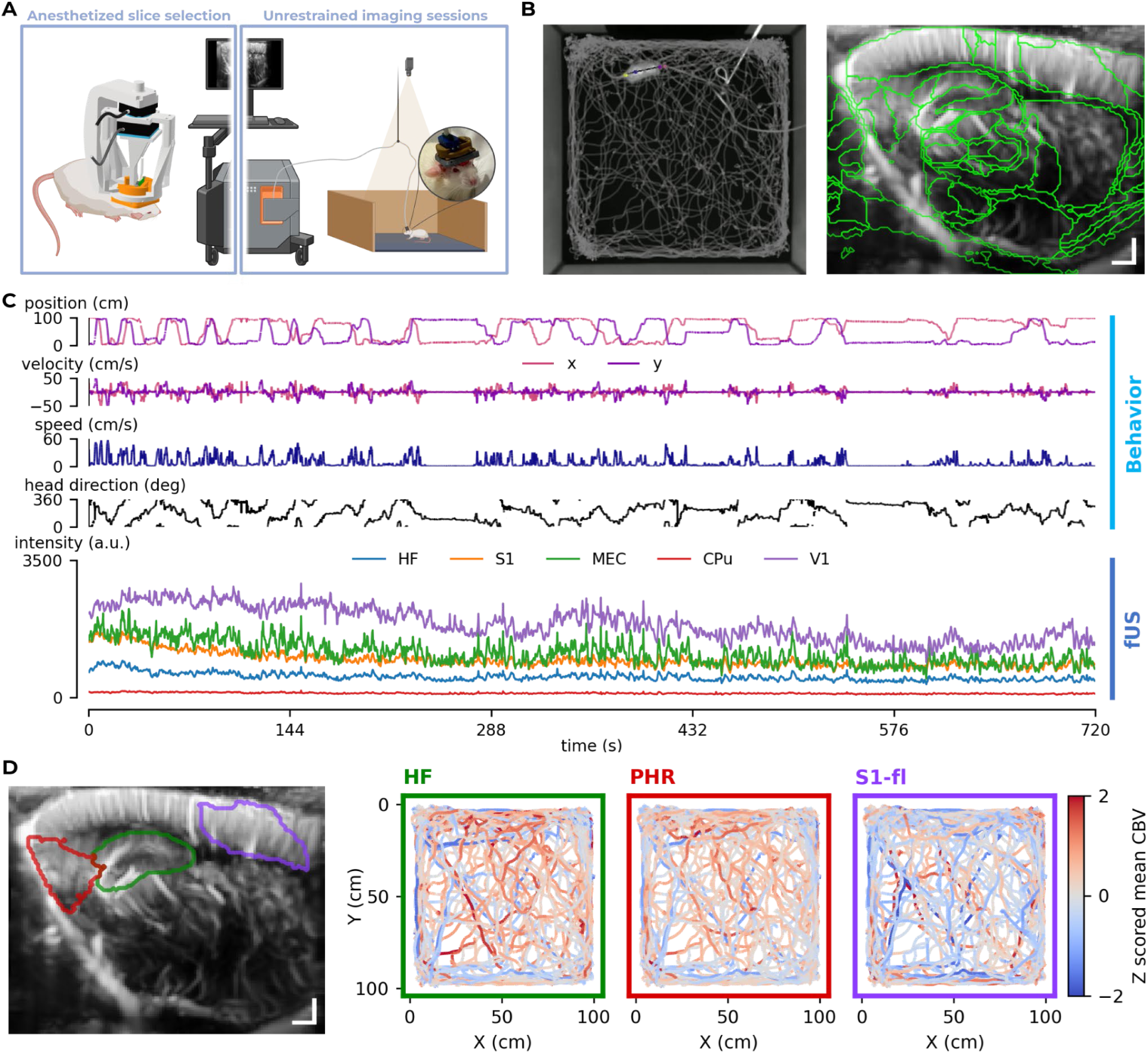
Fully integrated platform for fUS imaging of awake and freely-moving rats synchronized with behavioral signals to study the vascular dynamic response during spatial exploration: **A.** Scheme of the freely-moving functional ultrasound imaging experiments. Left, a single session of imaging is performed in each animal under anesthesia. This is necessary for the selection of the imaging plane and the calibration of the headset probe holder. Right: experiments in free exploration (in a 1 m2 open-field), the ultrasonic probe is plugged to the headset probe holder. **B.** Left, example of a video recording frame of an imaging session overlaid with the animal trajectory. Right, a typical imaging plane used in our study (3.8 mm para-sagittal) overlaid on the mean Doppler signal. **C.** Example in one recording of (non-exhaustive) behavioral signals extracted from the animal tracking data (from top to bottom, animal position on x and y axis, animal velocity on x and y, absolute animal speed, instantaneous animal head direction), and the concomitant region-wide averaged Power Doppler signals in the hippocampal formation (HF), primary somatosensory area (S1), medial entorhinal cortex (MEC), caudate putamen (CPu) and primary visual area (V1). **D.** Example ROIs represented on the imaging plane on the left (red: parahippocampal region (PHR), green: hippocampal formation (HF), purple: primary somatosensory area, forelimb representation (S1-Fl)) and the corresponding pre-processed Z scored mean CBV signal of these regions plotted as the colormap of the animal position in the experimental arena. In the Doppler frames, scale bars at the bottom right indicate 1 mm in each axis direction.

In order to investigate the correlation between CBV dynamics in the brain and spatial behavioral parameters, we used a lagged General Linear Model (GLM) approach. The lagged analysis consisted in building a series of design matrices in which the signal of interest (e.g. animal speed) was temporally shifted by one *dt* (*dt* = 80 *ms*) from a chosen range of lags [*t*_0_, *t*_1_[(Figure 2A). The lagged analysis allowed us to compute activation profiles for each voxel independently.

**Figure 2:**
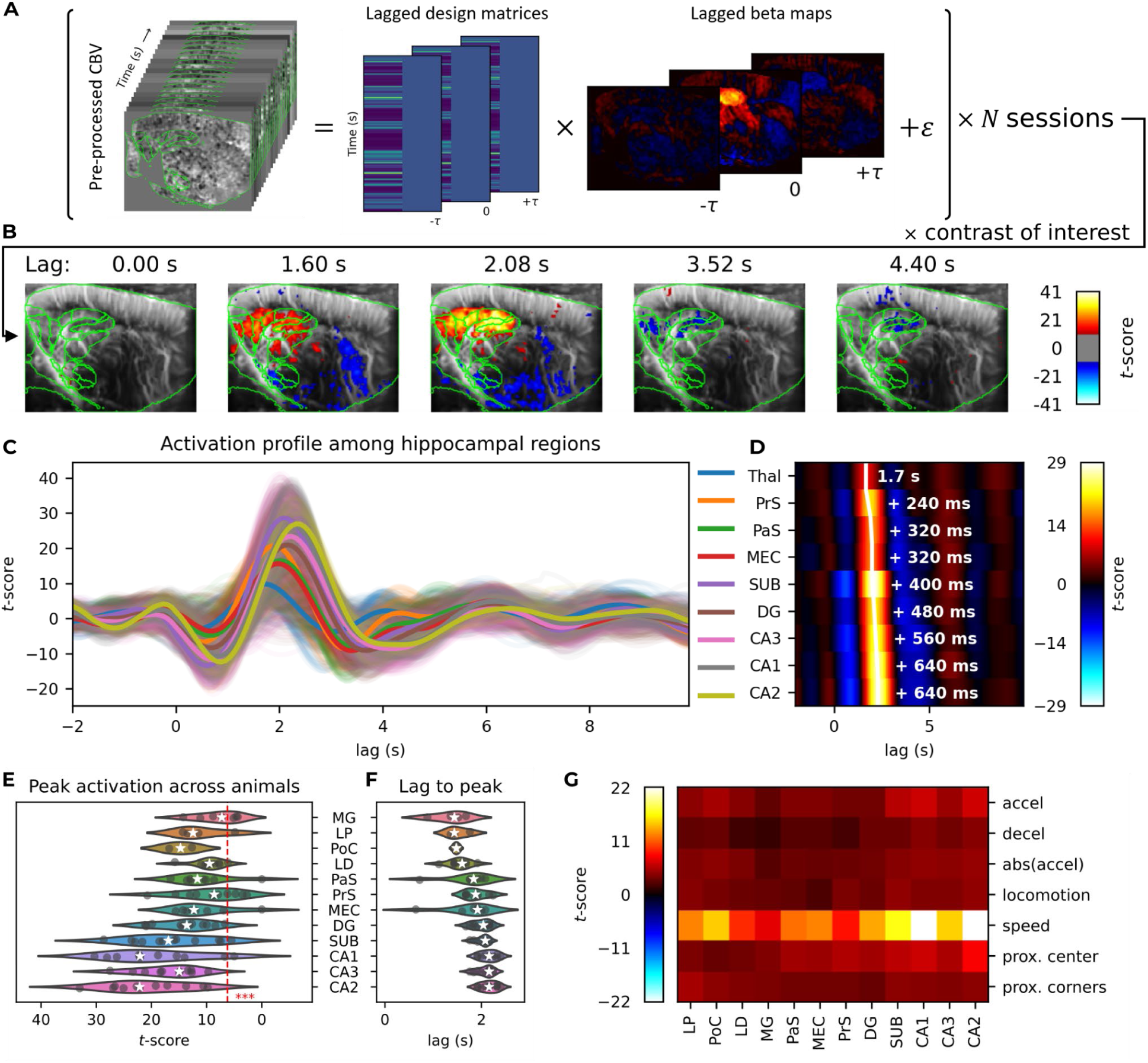
fUS imaging reveals robust correlation with animal speed during free exploration: **A.** Scheme of the GLM processing. Similar to a cross-correlation, one GLM is computed per lagged design matrix (lags from -2 s to 10 s). The result is a series of lagged beta maps, from which contrasts of interest are calculated to compute activation profiles. In this figure, the contrast is defined by the animal speed, i.e. the signal of interest. **B.** Examples of statistical maps at different lags from the animal speed contrast computed from the lagged GLM analysis proposed in **A**. Overlaid atlas regions in green represent ROIs from the hippocampal formation and dorsal thalamic regions. **C.** Activation profile from -2 s to 10 s of lag for an example animal. ROIs from the hippocampal region are plotted. Bold lines correspond to the mean value of the ROI, thin lines correspond to voxel-wise activation. **D.** Mean values for each region in **C** plotted as a heatmap. ROIs were sorted w.r.t the lag to peak in the activation profile. The lag to peak for each region is represented as the white line. The text to the right side of the white line for the top ROI shows the lag to peak for that ROI, while for the rest of the ROIs, the text shows the relative time difference between the lag to peak of the current ROI and the top ROI. **E.** Distribution of t-statistics value for the peak activation across imaging planes (7 animals, 12 imaging planes) in ROIs of hippocampal and thalamic regions present across imaging planes. **F.** Distribution of the lag to peak for the same ROIs across the same animals. ROIs were sorted according to the mean lag to peak. Red dashed line shows the t-score corresponding to the Bonferroni corrected p-value = 0.001. **G.** Heatmap showing median t-score values across imaging planes in the same ROIs as **F**, for different signals of interest that are somewhat correlated with the animal speed.

This analysis revealed robust activation in hippocampal and parahippocampal regions with respect to the animal speed. Figure 2A-C shows the analysis pipeline and an example where the lagged GLM of the animal speed was performed for 4 imaging sessions from lags of -2 s to 10 s. The resulting *t*-statistics maps (Figure 2B, Supplementary Video 1, Supplementary Figure 6) were computed from the session-averaged beta maps of the animal speed contrast and show specific activation patterns within the hippocampal formation. Different regions of interest in the hippocampal formation showed peak activation at different lag times, typically between 1.5 s and 2.5 s (Figure 2C-D). These peak latencies were consistent across animals and imaging planes, suggesting an directional flow of information from para-hippocampal areas in the medial entorhinal cortex (MEC, 1.92 +/-0.17 s), parasubiculum (PaS, 1.88 +/-0.13 s) and presubiculum (PrS, 1.92 +/-0.22 s), to hippocampal subregions including the dentate gyrus (DG, 2.04 +/-0.27 s) and cornu ammonis 1 (CA1, 2.16 +/-0.22 s), 2 (CA2, 2.16 +/-0.17 s) and 3 (CA3, 2.16 +/-0.16 s) (Figure 2F).

Furthermore, the lagged GLM analysis also revealed correlations, primarily in thalamic regions, when the animal head speed (i.e. the rate of change of the animal’s heading direction) was used as the signal of interest (Figure 3A-D), although these correlations were weaker than those observed with the animal speed. Figure 3A-D shows the statistical relationship between head angular speed and CBV changes in a representative animal using the same analysis framework as in Figure 2. Similarly, the activation peaked at a lag of approximately 2 seconds; however, the involved regions were now primarily located in distinct thalamic nuclei: the laterodorsal thalamic nuclei of the dorsal thalamus (LD), medial geniculate complex of the dorsal thalamus (MG), anterior nuclei of the dorsal thalamus (ANT) and posterior complex of the dorsal thalamus (PoC), and lateral posterior complex of the dorsal thalamus (LP). In some animals, weaker correlations were also observed in parahippocampal regions. However, since these thalamic structures are smaller than hippocampal and parahippocampal regions, they were not visible in all imaging planes, making it more challenging to confidently determine a possible sequential activation pattern underlying the information flow of angular head speed in our data.

**Figure 3:**
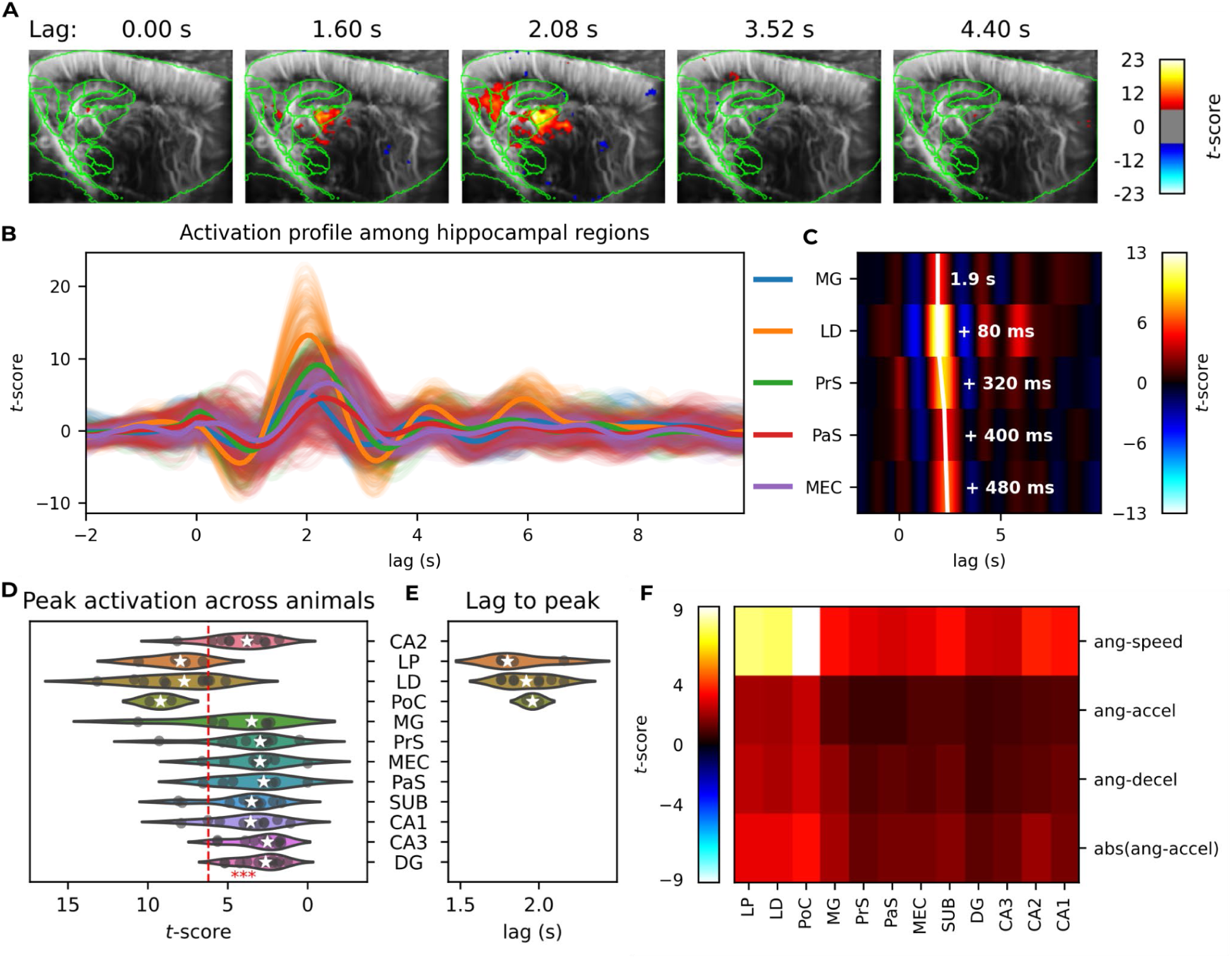
fUS imaging reveals correlation with angular head speed during free exploration: **A.** Example of statistical maps at different lags from the animal angular head speed contrast, computed from the lagged GLM analysis proposed in (Figure 2A). Overlaid atlas regions in green represent ROIs from the hippocampal formation and dorsal thalamic regions. **B.** Activation profile from -2 s to 10 s of lag for an example animal. Parahippocampal and thalamic ROIs are plotted. Bold lines correspond to the mean value of the ROI, thin lines correspond to voxel-wise activation. **C.** Mean values for each region in **B** plotted as a heatmap. ROIs were sorted w.r.t the lag to peak in the activation profile. The lag to peak for each region is represented as a white line. The text to the right side of the white line for the top ROI shows the lag to peak for that ROI, while for the rest of the ROIs, the text shows the relative time difference between the lag to peak of the current ROI and the top ROI. **D.** Distribution of t-statistics value for the peak activation across animals presenting the parahippocampal or thalamic region. Red dashed line shows the t-score corresponding to the Bonferroni corrected p-value = 0.001. **E.** Distribution of the lag to peak for the same ROIs across the same animals Only ROIs with median peak activation above 0.001 for a Bonferroni corrected p-value are shown. ROIs were sorted according to the mean lag to peak. **G.** Heatmap showing median t-score values across imaging planes in the same ROIs as **F**, for different signals of interest that are somewhat correlated with the animal angular head speed.

An exhaustive list of behavioral signals (including instantaneous speed, acceleration, head directions, place-like activity, grid-like activity, corner and wall proximity, angular head speed, and angular head acceleration) was evaluated using this lagged GLM approach. In most cases, we did not find any robust correlation between the CBV changes and these signals (Supplementary Figure 8). Some of these behavioral signals exhibited correlations among themselves at zero lag or with a small delay (for example speed and positive acceleration) and this can be seen in the pattern of activation responses in the same ROIs (Supplementary Figure 8). Although the spatial pattern of activations was similar across signals correlated with animal speed and angular head speed, the activation strength differed, with stronger responses observed for both animal speed and angular head speed (Figure 2G, Figure 3F, Supplementary Figure 8).

These results demonstrate that voxel-wise CBV dynamics in key regions of the spatial navigation circuitry are specifically and strongly associated with animal speed during free exploration. Although weaker than locomotion speed, head angular speed still elicited activation in thalamic regions across animals, further strengthening the link between local CBV changes and the rate of change in both position and direction during spatial navigation.

### fUS imaging reliably decodes locomotion speed and proximity to boundaries

To investigate how effectively CBV dynamics encode spatial parameters, we trained a set of decoders to predict animal speed, proximity to walls and proximity to corners. To account for the possibility that different voxels in our imaging plane may strongly correlate with the signal of interest, but at different time lags, we used the time-reversed lagged *t*-statistics values from the GLM analysis to convolve the time series of each CBV voxel and align them to an optimal phase (Figure 4A-D). The decoder was designed in two steps, performing first a linear principal component analysis (PCA) and then a ridge regression (for regression decoders) or logistic regression (for classification decoders). The penalty regularization parameter for the regressor/classifier and the number of PCA components were tuned in a grid-search style. After the convolution step, the data were split by session, and the ridge regression decoder was trained in a train-validation leave-one-session-out paradigm. Data from one session was held out as test data for the final evaluation of the decoders.

**Figure 4:**
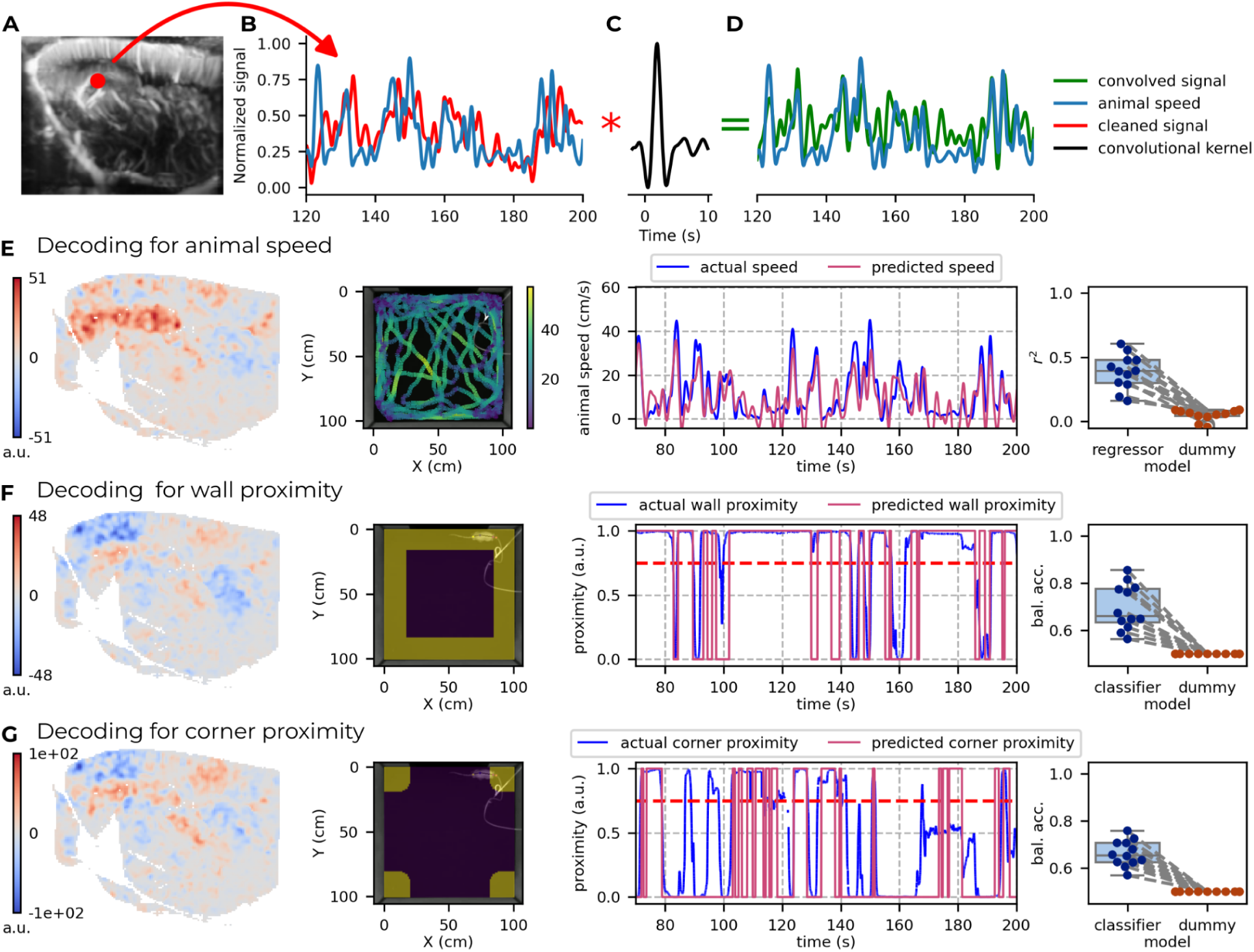
fUS imaging reliably decodes different spatial parameters for unseen sessions: **A.** Localisation of the voxel that exemplifies the convolution processing in **B**, **C** and **D,** done before the training paradigm. **B.** Pre-processed CBV time series of voxel in **A** (red) and animal speed (blue). **C.** Convolutional kernel used in this voxel taken from the GLM results (Figure 2). **D.** Convolved CBV time series of voxel in **A** (green) and animal speed (blue). **E, F and G.** Decoding output for animal speed, proximity from the walls and proximity from the corners, respectively. First column: Final weights output from each decoding pipeline. Second column: Visual insight of what information the decoder is searching for, i.e., animal speed (**E**), proximity from the walls (**F**) and proximity from the corners (**G**), respectively. Third column: Actual vs. predicted example plot for each decoding pipeline. The horizontal red dotted line in **F** and **G** shows where the threshold was set to define the two classes used in the decoder. Fourth column: Group level information represented by a box-and-whisker-plot where each dot represents one animal. Whiskers define the median, IQR and 1.5*IQR. All dots in the regressor/classifier column are directly compared to its dummy counterpart.

### fUS imaging decodes animal speed

The decoder trained to predict animal speed performed above chance in all animals and across all imaging planes, successfully generalizing to unseen sessions with a coefficient of determination of up to *r*^2^ = 0.6 relative to the actual speed signal (Figure 4E). The trained decoder consistently showed a preference for voxels in the hippocampal formation, indicating that CBV dynamics in this region robustly encode information about instantaneous animal speed (Figure 4E).

### fUS imaging decodes proximity from the walls and corners

Interestingly, for spatial parameters that did not exhibit robust activations in the univariate GLM approach, such as proximity to walls or corners, we were still able to reliably decode this information from CBV dynamics using the approach described above. The signal of interest was thresholded to create two binary classes (near the walls or not, and near the corners or not) and trained the decoder using a logistic regression to classify the final output instead. The decoders were also able to generalize well to unseen sessions, performing significantly above chance levels and achieving balanced accuracies of up to 0.85 for wall proximity and 0.8 for corner proximity (Figure 4F-G).

These results demonstrate that CBV dynamics within the spatial navigation circuitry can be used to reliably decode different spatial parameters during free exploration, even in cases where univariate approaches fail to reveal significant activations.

### Inter-animal decoding of the animal speed remains robust using region-wide fUS data

After successfully decoding animal speed in unseen sessions for a single animal, we aimed to test whether a decoder trained on one animal could predict the speed of a different animal using region-wide CBV dynamics. Instead of using voxel-wise time series as inputs at the beginning of our decoder pipeline, we extracted the mean CBV time series from anatomically defined regions consistently present across animals (CA1, CA2, CA3, DG, SUB, MEC, POR. PaS, PrS). This allowed us to incorporate animals imaged at different sagittal positions (for example, one at LR: -3.80 mm and another imaged at LR: +4.20 mm sagittal) within the same decoding pipeline (Figure 5B). The decoder was trained on a single animal and validated on 4 different animals. As expected, the results showed weaker performance compared to intra-animal decoders, but remained significantly above chance levels, with coefficients of determination reaching up to *r*^2^ = 0.36 relative to the true speed signal (Figure 5C). The decoding weights corroborated the strength of the activation patterns shown in Figure 2, demonstrating a strong preference for the hippocampal CA1 and CA2 regions, with slightly lower contribution from para-hippocampal regions (SUB, MEC and POR).

**Figure 5:**
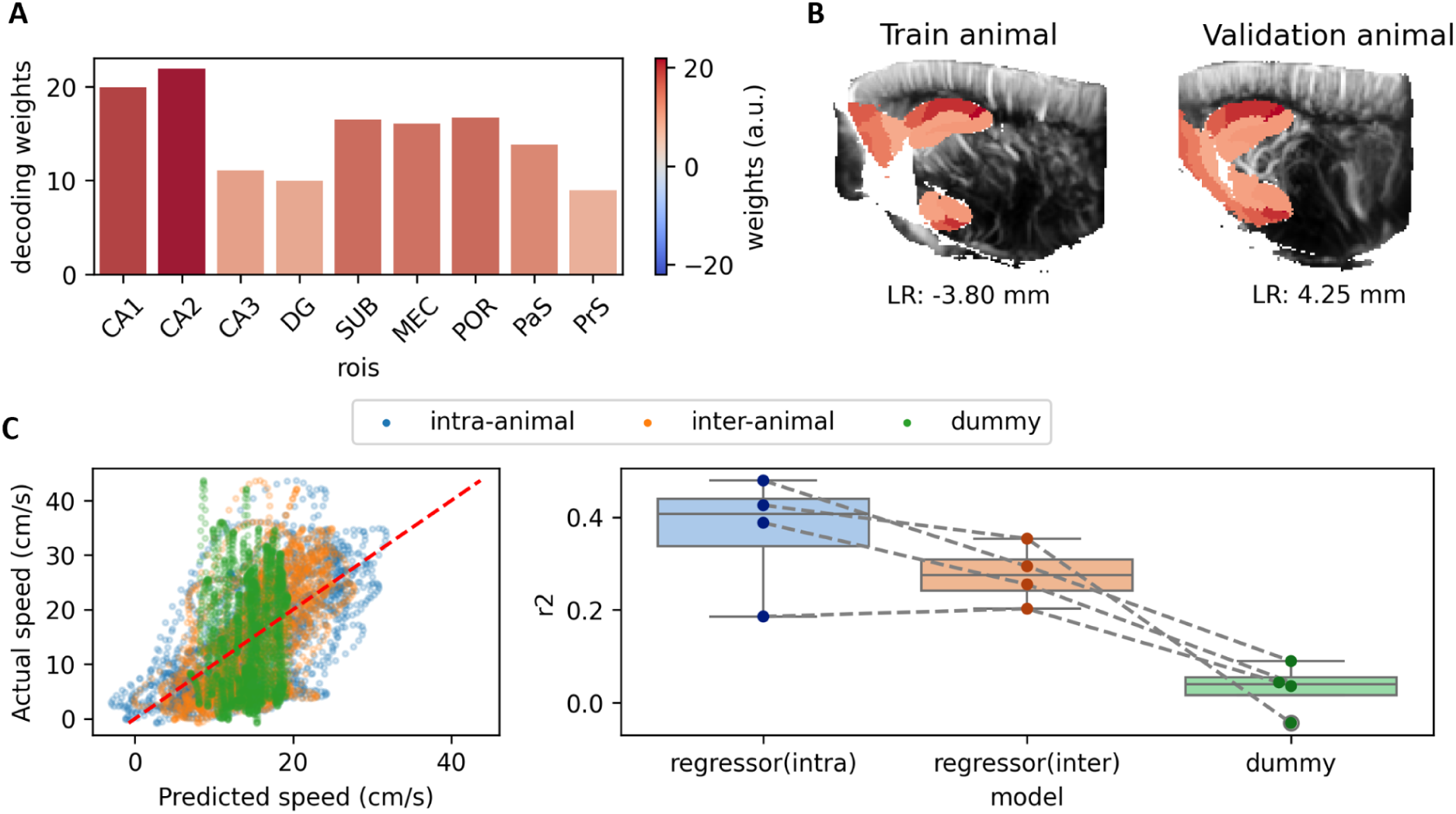
Inter-animal decoding of the animal speed using region-wide fUS imaging data: **A.** Decoding weights for each of the ROI used to train the inter-animal decoder (dorsal CA1, 2 and 3, DG, SUB, MEC, POR, PaS, PrS). **B.** The same weights as in **A**, but overlaid on the imaging plane of the animal used to train the decoder (‘Training animal’) and on one of the animals used as validation data (‘Validation animal’). The decoder was trained on a 3.80 mm left parasagittal plane (LR: -3.80 mm). The validation example shown in B was placed at the 4.25 mm (right) parasagittal plane. **C.** Left panel: Actual vs. predicted speed scatter plot containing the dummy decoder, inter-animal decoder and the intra-animal decoder (as in Figure 4). Right panel: Group level information on the same three decoders as in the left panel. Box-and-whisker plot where whiskers define the median, IQR and 1.5*IQR. Each dot represents one animal. For the inter-animal decoder, only one animal is used for training.

These results show that CBV-based encoding of animal speed is sufficiently robust to support decoding across different animals, even when the imaging planes differ and precise anatomical registration is not possible.

### Energy-speed scaling in a CAN model of path integration is in agreement with CBV dynamics

In order to understand how energy usage depends on animal speed, we examined a path integration system proposed in^52^. As illustrated in Figure 6B, the model features two ring attractors connected via both intra-ring and inter-ring couplings, which allow for forming bumps of neural activity pointing to the animal position and moving the bump according to self-motion cues. The bump profile is informative about the number and firing rates of the active place cells at any time. This model implements what is called a copy-and-offset double-ring network architecture, and is similar to the neural circuit sustaining the head-direction system^34^ of Drosophila^34^. The dependence of the energy rate on the animal velocity is quite robust to the model details, see below. Furthermore, while the two-ring model was originally introduced for path integration along a line, it can be extended to path integration in the exploration plane, without affecting the main results below.

**Figure 6:**
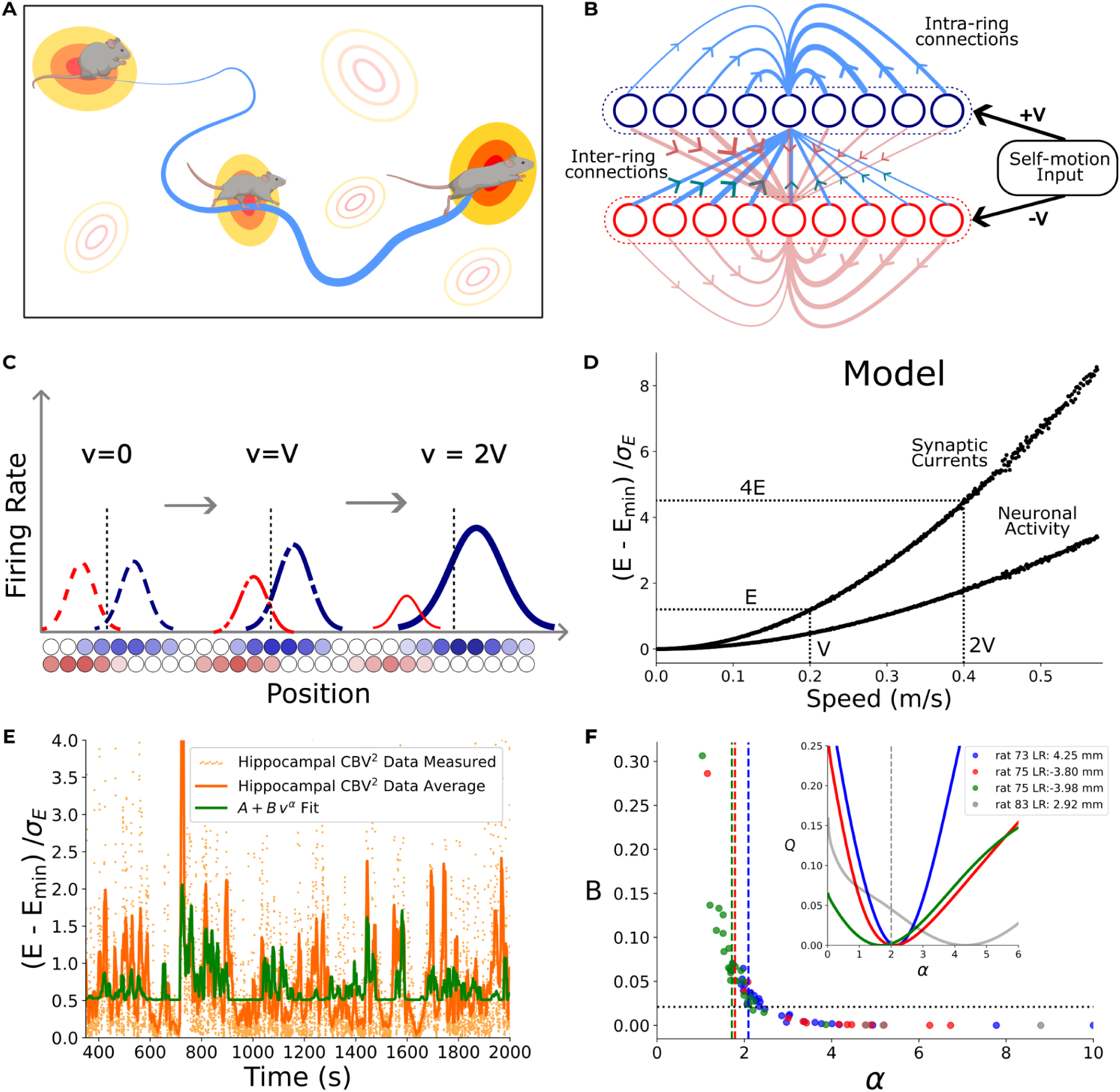
Path integration model predicts that energy consumption rate increases quadratically with velocity, in agreement with CBV measurements: **A.** As the animal explores the environment, the neural representation of its position is updated based on self-motion cues. This path integration process maintains the correspondence between the firing activity of place cells (indicated by color-coded firing maps) and the positions in space. **B.** Circuit for path integration based on the double ring model of Xie et al.^52^. Intra-ring recurrent connections allow for self-sustaining bumps to form. Inter-ring projections together with vestibular inputs allow for the two bumps to update their positions and carry out path integration. The width of the connections is indicative of their amplitude; the total activity is regulated by global inhibition (not shown). **C.** Bump profiles vary with the locomotion speed of the animal. At rest, the two bumps are identical. With increasing speed, the bump heights (defining peak neural firing rate) and widths (number of active place cells) change depending on the direction. Vertical lines locate the global centers of mass of the two bumps and show the encoded animal positions at different times. **D.** Energy rates in the double-ring model associated with synaptic currents and neural activity (with a 58/22 cost ratio between them^56^) vs. velocity, showing quadratic dependence. Path integration of the model was carried out on a 12 min experimental speed trace, with dots corresponding to time points. **E.** Hippocampal *CBV*^2^ time trace for animal 73 (LR: +4.25 mm) (orange scatter points: measurements, orange line: 10 s sliding average). The green line shows the result of the sliding average of the best fit *A* + *Bv*^α^, where *v* is the locomotion speed of the animal. **F.** Scatter plot of *B* vs. α for four slices (see color code). Each dot represents the result of the optimization procedure for a random group of sessions recorded for the same slice; groups for which A<0 were discarded. Vertical dashed lines show the average value of α for each slice. Inset: residual *Q* (after optimization over *A*, *B*) vs. α for two animals having large *r* ^2^ values in the decoding analysis. Rat 83 analysis led to small B and is shown in grey. (Figure 4E).

Informally speaking, the encoded position of the animal is defined as the average of the two bumps (Figure 6C). As velocity increases, the path integration mechanism results in one bump being wider and higher than the other and its position shifting at a rate proportional to the locomotion speed. Consequently, higher speeds amplify the disparity in bump sizes. Due to the approximate triangular shape of the bump, the increase in height and width, both proportional to velocity, leads to a quadratic increase in the largest bump area (Figure 6C). This qualitative property is largely insensitive to the details of the model connectivity and defining parameters.

As a result, mean neuronal activity quadratically increases with velocity. Similarly, the synaptic currents that support and shift the bumps over time show the same dependency (<u>Methods</u>). As energy expenditure increases linearly with action potential generation and the overall magnitude of postsynaptic currents, we expect a quadratic increase in energy consumption with velocity. This relation is confirmed by simulating path integration of a 12 min sample speed trace and calculating the energy consumed (Figure 6D). The quadratic dependence of energy on speed implies that small differences in speed at higher speeds result in disproportionately large differences in energy consumption.

We then compared this theoretical prediction with our experimental data. The data consists of fUS imaging (here called *BBBBBB* for simplicity) data from the hippocampus formation and the locomotion speed (*vv*) of the rat. The power relationship between CBV and energy consumption rate is approximately 2 in the literature (see <u>Methods</u>), so we take *BBBBBB*^2^ as a proxy for consumption. Inspired by our theoretical model, we look for the exponent αα in the equation *A* + *BBvv*^αα^ that best fits the *BBBBBB*^2^ dynamics. To avoid delay and distortion effects of the haemodynamic response function, we opt to minimize the differences between a sliding average of the CBV data with fixed window size τ = 10 s (see Supplementary Figure 9A), and the fitting function (Figure 6E). We proceed through least squares minimization (see <u>Methods</u>). After optimization over *AA*, *BB*, αα, we find that all slices for which a substantial correlation of CBV and velocity is found (large *B*), the exponent αα is close to 2 (Figure 6F), in very good agreement with the theoretical predictions. In addition, the sum of squared residuals, *QQ*(αα), is minimal for αα ≃ 2 for the animals and sessions that were top performers on decoding animal speed, see Figure 6F<u>, Inset and</u> Figure 4E). Cross-validation procedures and null models can be found in <u>Methods</u> and Supplementary Figure 9B,C. The best fit is shown against *BBBBBB*^2^ for one session in Figure 6E. While a clear correlation between the fUS imaging data and *vv*^2^ is observed, a large part of the time dependence of the CBV signal cannot be explained based on velocity alone. This is understandable considering the complexity and variety of factors affecting the haemodynamics in the brain^53–55^.

Lastly, we note that we also found evidence of quadratic scaling in the head-direction system in the thalamus. Although the evidence was less conclusive, due to the smaller size of the head-direction circuit and less angular velocity variation in the data, we were still able to see a correspondence for a couple of datasets (see Supplementary Figure 10C, D).

### fUS reveals cerebral mapping of minute-scale CBV oscillations during free exploration

The frequency content of the CBV signals is inherently slow and therefore very different from the frequency content of neuronal activity. However, recent studies in head-fixed rats have reported slow oscillatory neuronal activity in regions relevant to spatial navigation^57,58^. To investigate the presence of similar slow oscillatory patterns in CBV signals during free exploration, we calculated the autocorrelation of each voxel’s time series, followed by the corresponding power spectral density (PSD). The autocorrelation function of voxels within the same session (Figure 7A-B) reveals oscillatory patterns at consistent frequencies across specific brain regions. From the PSD of each voxel, we constructed PSD maps for each frequency bin. These maps often showed clusters of high spectral power in well-delineated brain regions, such as the hippocampus, visual cortex, somatosensory cortex and thalamic areas. Clusters delineating the hippocampal region were found in several sessions across different animals with peak frequencies ranging from 0.003 Hz up to 0.021 Hz (Figure 7C). To investigate if these slow oscillatory patterns could have a neurophysiological role in animal’s exploration, we also calculated the autocorrelation of the animal speed and its corresponding PSD. In 8 out of 10 sessions showing clusters of slow oscillatory patterns delineating the hippocampal region, the animal speed PSD exhibited a peak at similar frequency (Figure 7D). These results suggest that slow oscillatory patterns of CBV signals in the hippocampal region are related to the slow temporal dynamics of the animal’s exploratory behavior.

**Figure 7:**
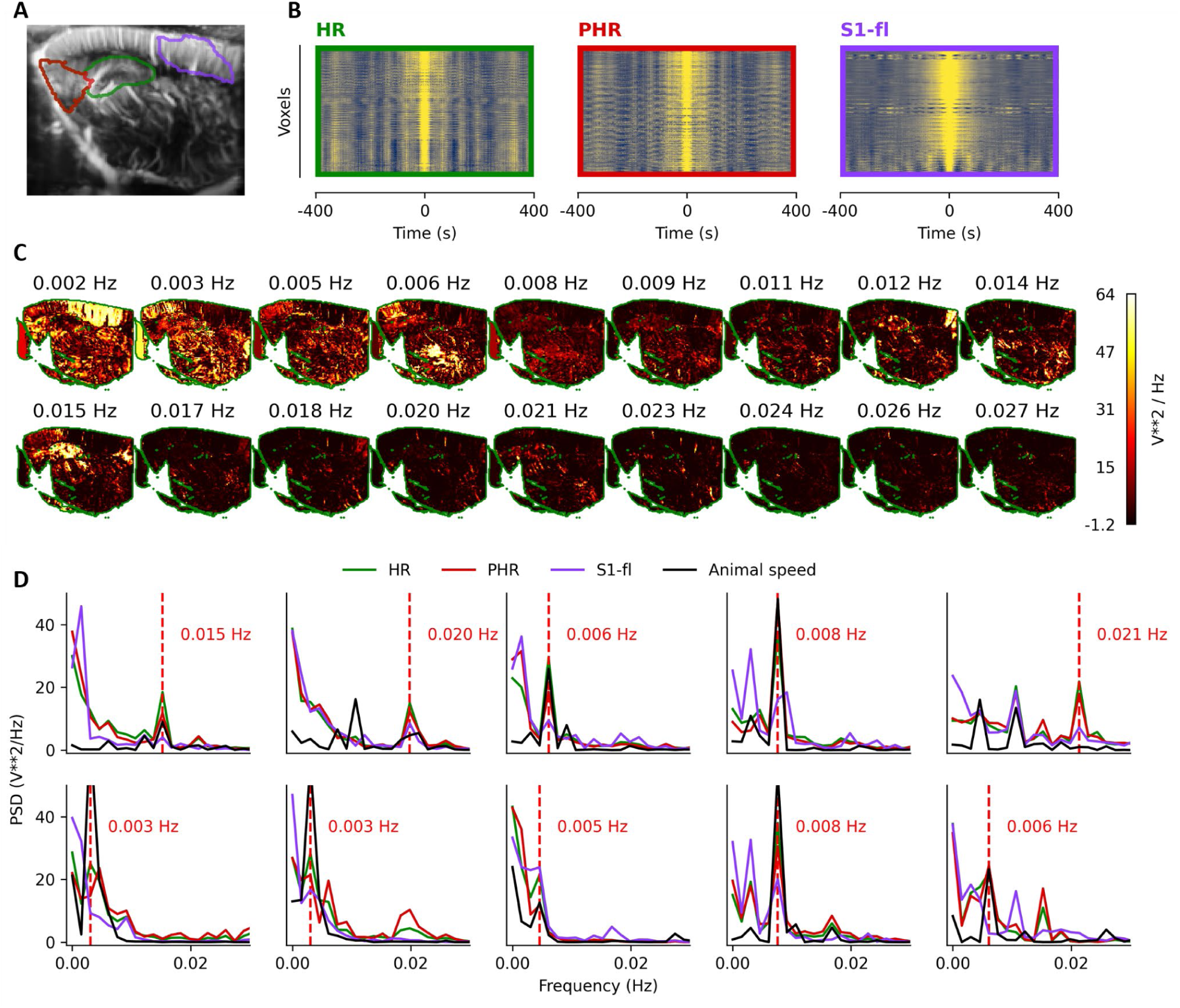
fUS reveals minute scale oscillation during free exploration: **A.** Power Doppler frame (same as in figure 1) with delineated ROIs. **B.** Carpet plot of the autocorrelation function of each voxel in the corresponding ROI (HR in green, PHR in red, S1-fl in purple) from -400 to 400 seconds of lag. **C.** Power spectrum intensity maps for a range of frequencies. Maps are constructed from the power spectrum density of the autocorrelation function of each voxel in the imaging plane. **D.** Power spectrum density in the example ROIs (HR in green, PHR in red, S1-fl in purple) and the power spectrum density of the autocorrelation function of the animal speed (in black). Vertical red dashed lines show the peak intensity frequency in the HR ROI. Each plot corresponds to a different animal or imaging session.

## DISCUSSION

In this study, we leveraged functional ultrasound (fUS) imaging to investigate the relationship between cerebral blood volume (CBV) dynamics and spatial navigation features in freely moving rats. Experiments performed during open-field exploration revealed that local vascular patterns within the hippocampal formation correlate strongly with locomotor speed, a critical variable to any navigation system. These findings provide novel insights into the neurovascular correlates of spatial navigation, suggesting that animal speed is encoded at the mesoscopic scale in the blood flow dynamics of the entorhinal-hippocampus circuitry.

Taking advantage of the high spatiotemporal resolution of fUS imaging data (100 µm, 80 ms), we found that local blood flow dynamics within the hippocampal formation robustly correlated with instantaneous animal speed. The same pattern of activation was consistently found across 7 animals, and 12 different imaging planes (Supplementary Video 1, Supplementary Figure 6). Notably, the temporal structure of this correlation revealed a sequential activation progressing from thalamic nuclei to parahippocampal regions and ultimately to the hippocampus, consistent with the hierarchical processing of spatial information in the navigation circuit. The observed delays in peak activation (∼1.5-2.5 s) align with previously reported hemodynamic response function (HRF) latencies in rodents^42,46,47,59^, further validating the neurovascular coupling captured in our fUS imaging measurements.

FUS imaging reveals a sequence of activation from the thalamus to the medial entorhinal cortex (MEC), then to the dentate gyrus, and subsequently to the CA subregions of the hippocampus. This spatiotemporal progression is consistent with previous electrophysiological studies performed using EEG recordings in rats, which have similarly highlighted the directional flow of information from the entorhinal cortex to the hippocampus during navigation^3,60^. Theta rhythms, a hallmark of spatial navigation, exhibit a coordinated phase relationship between MEC and hippocampal regions, with MEC theta activity often leading hippocampal activity^3,61^. This phase lead is thought to support the integration of self-motion cues and environmental inputs, facilitating the dynamic updating of spatial representations in the hippocampus. Moreover, the stronger and earlier activation observed in the MEC aligns with its well-established role in path integration and grid cell-based spatial encoding, which provides a metric framework for navigation^1^. The subsequent activation of hippocampal regions likely reflects the integration of positional and directional information to generate place-specific representations, as observed in place cell activity during navigation.

Furthermore, our data also show a correlation between angular head speed and several nuclei in the dorsal thalamus, regions known to contain a high proportion of head direction cell population receiving vestibular inputs from regions containing angular head velocity cells^9,62–67^. Suggesting that this vascular demand supports the transformation of angular velocity encoding into angular heading direction happening in these dorsal thalamic regions^65,66^.

The lack of clear vascular signatures for place cell-, grid cell-, or head direction cell-like patterns is likely due to the high-dimensional and distributed nature of these spatial representations. Encoding *N* different place fields or grid modules requires *N* distinct neurons, and these representations lack a defined anatomical clustering. As a result, any given fUS imaging voxel covering hundreds to thousands of neurons, inevitably averages signals from neurons tuned to different spatial codes. Two factors make this particularly challenging for fUS imaging: (1) spatially-tuned neurons (e.g., place or grid cells) do not present any specific spatial arrangement regarding their location in the brain in a way that would allow spatial summation of their signals at the mesoscopic scale; thus, voxel-averaged CBV changes dilute meaningful activity patterns. (2) only a subset of these spatially-tuned cells is active at any given time, since the animal cannot occupy multiple places or face multiple directions simultaneously. Yet, during exploration, the animal is always somewhere and heading in some direction, resulting in a temporally continuous but spatially scattered activity across the whole hippocampal formation during locomotion. From the fUS imaging perspective, this manifests as broad and distributed CBV changes rather than localized signatures, making it difficult to resolve individual spatial codes.

On the other hand, speed encoding is unidimensional-each speed cell conveys the same type of information, with firing rate increasing proportionally with locomotor speed. This characteristic is particularly advantageous for fUS imaging, as the entire population of speed-encoding cells will generate a synchronized vascular demand at the same temporal scale. Interestingly, Gois et al.^20^ demonstrated that speed cells in CA1 are primarily fast-spiking inhibitory interneurons, exhibiting firing rates that can be several times higher than those of pyramidal or stellate cells, which typically constitute the population of classical spatially-tuned cells^68–70^. Similarly, angular head speed signal may offer the same advantage for fUS imaging as locomotion speed, as it can also be encoded unidimensionally, potentially leading to synchronous and linear vascular demands. Even though the exact linearity of the neurovascular coupling across fast-versus slow-spiking neurons is yet to be elucidated, it is plausible that a fast-spiking neurons population will generate a higher metabolic demand and therefore larger neurovascular response than other neurons for the same stimulus^71,72^. Speed-related vascular dynamics may reflect metabolic demands associated with sustained locomotion or vascular coupling to speed-modulated neuronal activity, such as hippocampal theta oscillations. Additionally, they could correspond to the metabolic demand of fast-spiking interneurons involved in gamma oscillations, which are known to play a key role in path integration and communication between the MEC and hippocampus^71,72^. This raises intriguing questions about whether vascular changes in these regions might contribute directly to navigation mechanisms, rather than merely serving as indirect proxies for neuronal activity.

Additionally, our study demonstrates that fUS imaging data can be used to reliably decode animal speed in unseen imaging sessions, achieving coefficients of determination up to 0.6 in certain imaging planes. Interestingly, similar decoding performance has been reported when using neuronal activity from speed cells in the MEC and hippocampus^18,20^. The ability to train a decoder model once and apply it to newly acquired data without retraining significantly reduces the need for frequent recalibration, reinforcing the longitudinal stability and reliability of the hemodynamic signals captured by fUS imaging. Furthermore, despite variability in imaging planes across animals, we successfully trained above-chance level decoders for animal speed using ROI-averaged CBV signals, suggesting that local blood flow dynamics encode speed in a stable and reproducible manner across animals, paving the way for more generalized modelling of behavioral parameters in future studies.

To better understand why and how the animal velocity is related to CBV data, we have developed a theoretical approach based on a continuous attractor network (CAN) for space representations in the rodent brain. In CANs, a collective bump of activity of spatially-tuned cells encoding nearby representations in space is updated from sensory inputs as the animal moves. Proprioceptive inputs alone, carrying information about speed (and acceleration) is the basis for path integration, that is, the capability to accurately update the neural representation of the position. While models for path integration in CAN were introduced two decades ago, first for angular orientation, then for position, their energetic implications have not been investigated so far to our knowledge. Revisiting classical CAN models^52^ with a proxy for energy consumption taking into account the production of spikes and synaptic currents, we predict that the energy rate increases with velocity, with an exponent α = 2. The reason for this quadratic increase is that the path integration mechanism changes both the height and the width of the bump linearly with speed, leading to a quadratic effect on the bump area, and thus on the mean firing activity and on the mean currents. As a result, the value α = 2 is quite generic and does not depend on the particular implementation of the network. Alternative models for one-dimensional CAN developed in the past would also show a similar dependence^73–76^.

To confirm this prediction, we have analyzed the CBV data in the hippocampal formation from the fUS imaging recordings and fitted its time behavior against the speed traces of the locomotion of the rats. We find, for all sessions in which a substantial correlation exists, that the best fit is (close to) quadratic. This result provides indirect evidence for the presence of CAN-supported path integration in hippocampal representations. We emphasize that this analysis does not aim at explaining the entire CBV signal, but only its dependence on velocity. A strong time-dependent residual can be observed in the fitted model.

While α = 2 is a robust prediction of most theoretical models, it is not unique. For instance, models in which the bump height varies with velocity more than the width would lead to α comprising between 1 and 2. This scaling is seen in a computational model for Drosophila, where the active neural population is limited in size and show small variation with speed, see Supplementary Figure 10A,B. Generally, the value of α is informative about how the bump is reshaped by velocity changes. This can be directly characterized by electrophysiology experiments. Several investigations have found a linear relationship between place-cell activity and speed^20,77,78^. To our knowledge, no experimental evidence for the change in width of the bump has been reported; however, it is experimentally more delicate to quantify the size of the bump than its height, as it requires a precise comparison with background activity of the entire hippocampal neural population.

Extracellular firing-rate recordings capture only action potentials, but the bulk of the brain’s metabolic cost arises from synaptic currents^56^, which are notoriously difficult to measure directly via electrophysiology; fUS imaging, through neurovascular coupling, integrates both spiking-and synaptic-driven demands, providing a more comprehensive window into neural energy use^47,48,55,79,80^. Our recordings were made without controlling for visual feedback and repeating them in complete darkness -or ideally in head-fixed VR setup, eliminating vestibular inputs and allowing precise manipulation of locomotor gain -would help isolate the contribution of pure self-motion signals to the observed CBV-speed scaling. We note that previous work points to place cells maintaining their speed-modulated firing regardless of optic-flow manipulations^81^.

We also identified slow, minute-scale oscillatory patterns in CBV signals within the hippocampal formation. Previously evidenced using calcium imaging and electrophysiology^82,83^, similar oscillations have been linked to temporally ordered external landmarks even in environments limited to tactile and visual cues. These oscillations, occurring at minute-scale frequencies, may reflect coordinated vascular and neural dynamics underlying exploratory behavior. Interestingly, in the absence of external cues (exteroception), recent studies have reported similar slow oscillations in the medial entorhinal cortex^57^ using both calcium imaging and neuropixels recordings, suggesting that these rhythms might also play a role in maintaining temporal coordination across brain regions during spatial navigation. Furthermore, in the absence of external memory demand or spatiotemporal boundary, internally recurrent sequences were also found in the hippocampus as a template of spatiotemporal information during self-motion navigation^58^. Although our setup stands as a minimalistic free exploration setup (square arena without any explicitly designed exteroceptive cues, rewards or other explicit sensory cues), exteroception cannot be eliminated. Animals inevitably process subtle visual, tactile, olfactory, and auditory cues from the experimental environment. However, the absence of explicit exteroceptive cues in our experimental setup provides here a self-motion-based navigation, where proprioceptive and vestibular signals dominate. In this context, we show that hippocampal CBV oscillations at minute-scale mirrors oscillations present in the animal speed signal. Interestingly, fUS imaging provides brain-wide spatial mapping of these oscillations nicely delineating different brain regions (hippocampal formation, sensory, motor and visual cortex) with different oscillatory frequencies. In light of these results, the possible presence of minute-scale oscillations in the animal speed from former works^57,58^ would be interesting to investigate.

By leveraging fUS imaging to study spatial navigation in freely exploring rats, this study provides novel insights into the mesoscopic vascular correlates of navigation-related parameters, particularly animal speed. Our findings complement traditional electrophysiological approaches, offering a broader perspective on how spatial navigation information is represented at the level of cerebral blood volume dynamics. Our decoding analyses further demonstrate the potential of CBV signals to encode spatial parameters and the ability to decode animal speed across sessions. Furthermore, decoding across animals highlights the robustness of fUS imaging and the universal nature of this encoding. Although voxel-wise GLM analysis was found to be much less sensitive to features such as boundary and corner proximity, our ability to successfully decode these signals underscore the utility of multi-voxel population-level patterns in revealing subtle behavioral correlates that are not immediately evident in single-voxel analyses.

Still, our ultrasound images capture a narrow volume of the rat’s brain, not allowing for synchronous analysis of the entire spatial navigation system, but volumetric ultrasound imaging is a fast developing research field and we might be able to do 3D fUS imaging in freely moving rats sooner rather than later. Future work could also combine fUS imaging with electrophysiology or optical imaging to link vascular signals more directly to underlying cellular activity, and explore whether mesoscopic CBV dynamics reflect not only overt behavior but also cognitive processes such as planning, memory, or decision-making during navigation.

Our findings extend the understanding of spatial navigation to the vascular domain, demonstrating that CBV changes follow a similar temporal sequence, potentially reflecting underlying neuronal activity patterns. These results also underscore the capacity of fUS imaging to resolve the chronology of brain dynamics with a temporal accuracy well below the scale of neurovascular coupling delays and cardiac pulsatility. Together, these findings suggest that the spatial navigation circuit integrates speed information in a temporally and spatially organized manner, supporting a dynamic framework for path integration and spatial memory.

## METHODS

### Anim als and experiment al design

All experiments performed in this study complied with the French and European Community Council Directive of September 22 (2010/63/UE). Seven male Sprague Dawley rats were used with the approval of the local Institutional Animal Care and Ethics Committees (#59, ‘Paris Centre et Sud’ project #19701 2019031020578789 V5). Accordingly, the number of animals in our study was kept to the minimum necessary. To do so, animals were imaged several times and at different imaging planes, resulting in 12 different imaging planes and 58 imaging sessions (see Table 1). One week after their arrival in the laboratory (aged 8 weeks), the animals were housed by two and given a week of habituation with the experimenter and then underwent surgical craniotomy and implantation of an ultrasound-clear prosthesis. After the surgery, the animals benefited from a one-week recovery before getting habituated to the experimental setup, ultrasonic probe and ultrasonic probe holder. All rats were fed *ad libitum* and housed on a 12-h light/dark cycle in cages of two after chronic window implantation.

**Table 1:**
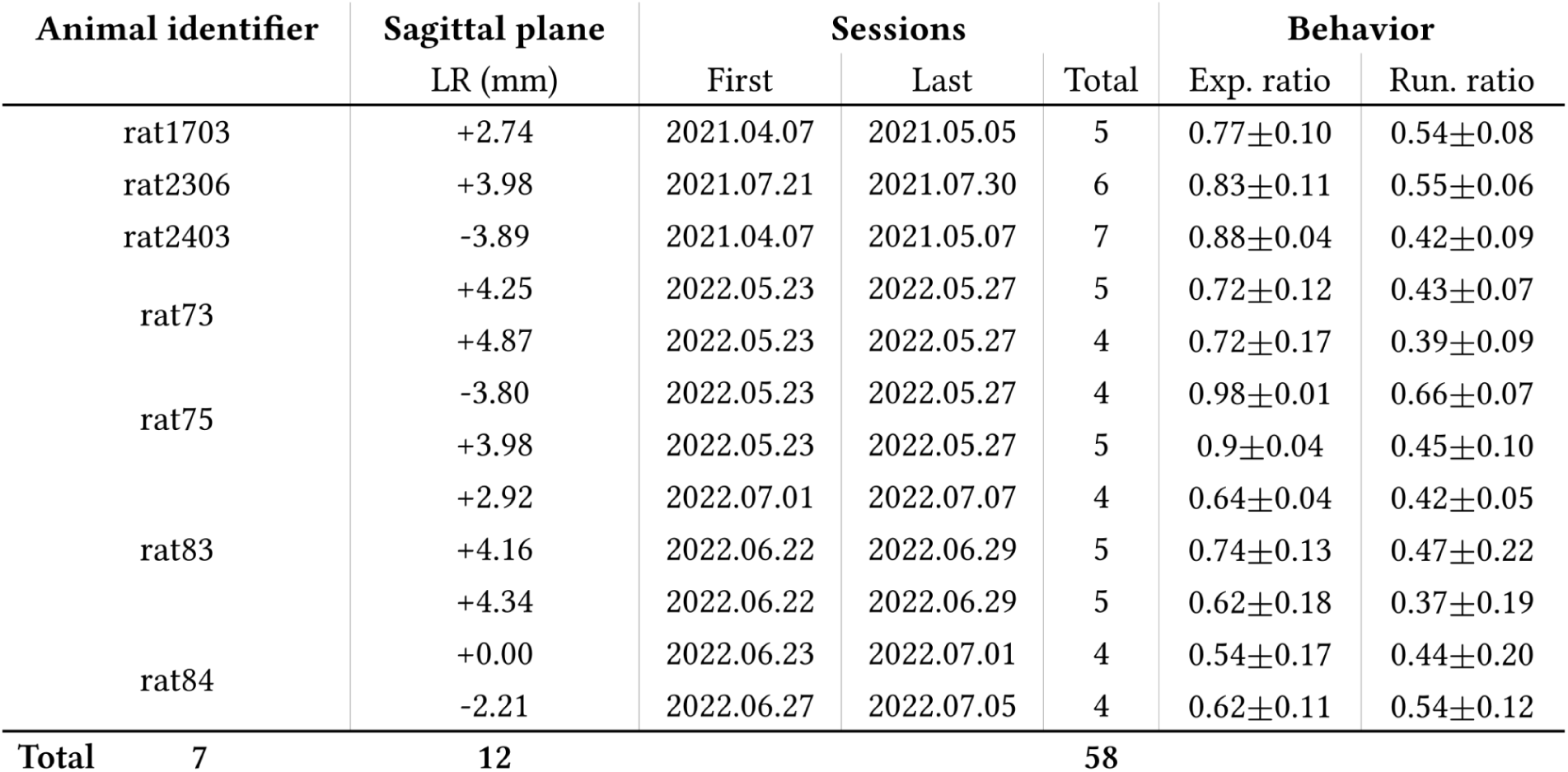
Extensive list of fUS imaging experiments performed on freely-moving rats: Parasagittal planes are identified by the left-right (LR) distance from the midline (negative to the left and positive to the right). Within sessions, dates of first and last sessions are given. For the behavior columns, data is given as mean +-std across sessions. <u>Exp. ratio:</u> Exploration ratio (ratio of visited bins, 10×10 binning of the arena). <u>Run. ratio:</u> Running ratio (running time over all time).

To position the probe on the desired imaging field of view, in a single anesthetized session of imaging, the brain vasculature was scanned, the plane chosen and the probe holder fixed for this plane (see Supplementary Figure 1 for the probe holder pieces). Experiments in freely moving animals then took place every other day (see below for further details) for up to 5 weeks. After the final experiment, animals were sacrificed using an overdose of pentobarbital and an intracardiac perfusion of 4% paraformaldehyde.

### Implantation of an acoustically transparent chronic window

As previously described^43^, following an initial induction of the anesthesia (4% isoflurane in 70% air, 30% oxygen), the anesthesia was maintained using 1,5% isoflurane, subcutaneous injection of buprenorphine 0,05 mg/kg 20 min before surgery and anesthetized locally with Lurocaine 4 mg/kg while body temperature was maintained at 37.0 °C with a heating pad (Phymep, Paris, France). Respiration and heart rate were monitored using a PowerLab data acquisition system with the LabChart software (ADInstruments, USA). The eyes of the rat were protected using an ointment (Ocry-gel, TVM, UK). A sagittal skin incision was performed across the posterior part of the head to expose the skull. We excised the parietal and frontal flaps by drilling and gently moving the bone away from the dura mater. The opening exposed the brain from Bregma +4.0 to Bregma −9.0 mm, with a maximal width of 14 mm. A prosthetic skull made of a ultrasound transparent material (polymethylpentene (=TPX), 250 μm thick thermoformed on the brain’s shape (Goodfellow, Huntington UK, goodfellow.com)) was sealed in place with Superbond C&B (Sun Medical, USA) and Tetric Evoflow IVOCLAR VIVADENT(Henry Schein, France), and the residual space was filled with saline. We chose a prosthesis approach that offers a larger field of view and prolonged imaging condition over 4–6 weeks compared to the thinned bone approach^42,43^. This material has tissue-like acoustic impedance that allows undistorted propagation of ultrasound waves at the acoustic gel-prosthesis and prosthesis-saline interfaces. Particular care was taken not to tear the dura to prevent cerebral damage. The surgical procedure typically took 4-5h. Animals were subcutaneously injected with anti-inflammatory drug (Metacam 2 mg/kg) and prophylactic antibiotics (Borgal, 30 mg/kg), and postoperative care was performed as long as experiments were being performed. Animals recovered quickly and were used for data acquisition after a conservative two-week resting period.

### Selection of the imaging plane and probe positioning

As we wanted to image reproducibly in the same imaging plane throughout our experiments, the optimal imaging plane had to be identified in each animal and in order to allow a fast, easy and reproducible positioning of the probe in the probe holder the determination of this plane and of the probe holder were determined and fixed in a single imaging session performed in anesthetized animals (Figure 1A). For each animal we selected between one or three brain-wide sagittal imaging planes covering the hippocampal and parahippocampal regions, due to their particular relevance for spatial navigatio^1^.

Animals were anesthetized using isoflurane (induction 4% in 70% air, 30% oxygen, followed by 1,5% after 10 min). They were placed on a stereotaxic frame and kept at 37°C using a heating pad (Physiotemp). The eyes were protected using an ointment (Ocry-gel, TVM, UK). A custom 2 axis motor could control the 2 degrees of freedom of the probe holder (rotation around the z-axis and translation orthogonal to the rotation axis) to place the probe in the desired imaging plane (Figure 1A).

The custom made 3D-printed probe holder was attached to its matching piece on the animal’s skull (Figure 1A). Probe positioning within the probe holder was performed either manually or using an automatized motorized setup (Supplementary Figure 1) based on 3D brain atlas registration during calibration experiments dedicated to probe holder optimization.

### Functional ultrasound (fUS) imaging

#### fUS imaging in freely exploring rats

At the beginning of each recording session, while the animal was awake and unrestrained in the arms of the experimenter, 1 mL of acoustic gel was applied to the skull prosthesis, the ultrasound probe (IcoPrime lite, 15 MHz, 128 elements, 0.110 mm pitch, Iconeus, Paris, France) was positioned in the ultrasound probe holder and both were attached to the animal’s head using custom 3D printed pieces (Supplementary Figure 1). The animal was then placed individually in a custom-made squared arena (1 m^2^, with 50 cm high walls) and the exploration started immediately, just one or two seconds before the beginning of the functional ultrasound imaging recording. Every recording session lasted for 12 minutes. The same animal was recorded at least 4 times in different days with different imaging gaps between sessions/animals.

#### fUS sequence

We performed multiplane wave functional ultrasound imaging using an ultrafast ultrasound acquisition system (Iconeus One, Iconeus, Paris, France). Each insonification of the medium was done using 8 successive plane waves with the following angles (−7°, −5°, −3°, −1°, 1°, 3°, 5°, 7°)^36,84,85^ at 4000 Hz pulse repetition frequency (PRF). As previously described^85^, the summation technique results in multiplane wave compounded frames at 500 Hz, that is, a coherent summation of beamformed complex in phase/quadrature (IQ) images. Data was acquired continuously. An SVD-based clutter filtering^86^ was used in each block to differentiate and separate tissue from blood motion and thus obtain a Doppler image (see *<u>Doppler image computation</u>*).

#### Anatomical atlas registration

For each sagittal recording, the imaging plane was registered to sagittal sections of the Waxholm Space (WHS) Atlas of the Sprague Dawley rat brain^87^ using anatomical landmarks as references, such as the hippocampus, cortical regions and large blood vessels. Minimal scaling and rotation was performed along each axis (Figure 1B, Supplementary Figure 2).

#### Automatic Doppler registration

To analyze data from the same imaging plane across days, we registered each imaging session to a reference session in a cross-validated manner. We used custom Python code leveraging SimpleITK registration utils and methods^88^. A global correlation metric was retained for each imaging session and the reference session. The correlation metric was calculated using the power Doppler frame (in decibels) of an imaging session that presented the lowest sum of intensity. We kept the reference and associated sessions that output the best average correlation metric. This was performed for each imaging plane (Supplementary Figure 3).

### Signal analysis

Data pre-processing.

#### Doppler image computation

With the WHS atlas correctly registered to an imaging plane, we re-computed the Doppler images from the consecutive blocks of 200 IQ images by using the SVD-based clutter filtering only on brain voxels. We removed the first 60 SVD modes and integrated the power of the rest of the modes in blocks of 200/5 = 40 IQ images, resulting in Doppler images at 12.5 Hz.

#### Framewise registration

For each imaging session we performed framewise registration using custom Python code leveraging SimpleITK registration utils and methods^88^. As reference, we used the imaging frame presenting the minimum overall sum of voxel intensity.

#### DVARS outlier detection

DVARS metric^89,90^ was calculated per imaging session. Voxels in imaging frames presenting relative DVARS above 4 were replaced by their temporal mean to a maximum of 4% of the total imaging frames in an acquisition session.

#### Temporal filtering

For each imaging session we performed butterworth (3rd order) band-pass filtering from 0.01 to 0.5 Hz.

#### White matter regression

For each acquisition session, we regressed the white matter averaged time series from each voxel in the brain via QR-decomposition

### Lagged GLM

A General Linear Model approach was adapted from fMRI Python packages^91–93^ to work on fUS imaging data. Instead of convolving the signal of interest in the design matrix of the GLM approach with different HRFs, we chose to perform a lagged approach. A GLM calculation was done across sessions for each imaging plane, for a range of lags applied to the signal of interest (typically from -2 to 10 seconds), resulting in one beta map per lag. Statistical parametric mapping^94^ could then be carried out to output a lagged series of t-statistics maps.

### Decoding

#### Preprocessing of the signal of interest

Before the training step, we preprocessed the output of interest in the same way we preprocessed the imaging data (see <u>Data pre-processing</u>), i.e. temporal filtering and white matter regression, to match both data distribution in terms of frequency content.

#### Preprocessing of the training data

We performed the Lagged GLM in the training subset against the signal of interest. As a result, we have a “GLM response function”, similar to a cross-correlation function, for each voxel. Since the optimum maximum for each voxel is possibly different in lag, we used the time-reversal of the GLM response function for each voxel to convolve the preprocessed CBV data itself, thus using the lag information and the strength of the activation available from the Lagged GLM analysis.

#### Cross-validation strategy

For every intra-animal decoding analysis, data was divided into training and test sets in a leave-one-session-out strategy. The decoder further divided the sessions within the training set in training and validation sets, again in a leave-one-session-out strategy.

#### Training pipeline

We used *scikit-learn*^92,93^ pipelines to build our training paradigm. The pipeline was composed of a Principal Component Analysis (PCA) step and of a final ridge regression step.

### Minute-scale oscillations

#### Autocorrelations and voxel-wise spectral analysis

We applied a similar method to Gonzalo Cogno et al.^57^, adapting it to voxel-wise time-series. For each voxel inside the brain mask, the PSD was calculated on the autocorrelation of its CBV fluctuations. The PSD was calculated using the scikit-learn implementation of the FFT. The PSD was then averaged within atlas-based regions of interest. A peak-detecting algorithm starting from 0.002 Hz was then used in these averaged PSDs.

### Cont inuous At t ract or Net work Model

#### Two-ring model for path integration

The dynamical equations for the firing rates *f* of neurons coding for position *x* at time *t* are given in Xie et al.^52^ and read, using − and + subscripts to denote the two rings *f*_±_(*x, t*) =

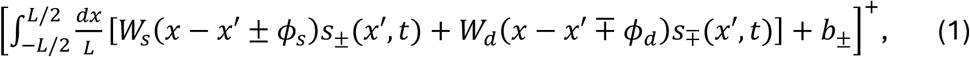

where [⋅]^+^ = *max*(0,·), and the vestibular inputs that differentially signal movements are denoted by *b*_±_ = *b*_0_ ± *v*, where *vv* is the velocity. Assuming periodic boundary conditions, the intra- and inter-ring connectivity matrices are given by, respectively,

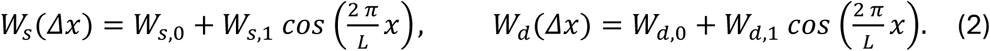

The parametrization will be fixed (unless stated) to *W*_*s*,0_ = −60, *W*_*s*,1_ = *W*_*d*,1_ = −80, *W*_*d*,0_ = −5. The inter and intra-ring asymmetry phases are equal to, respectively, 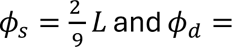 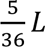 (see Xie et al.^52^ for details on the effects of different parameter choices).

Variables *s*_±_represent synaptic activation in each ring (i.e. the neurotransmitter concentration in the post-synapse) and it is determined by a set of first-order differential equations

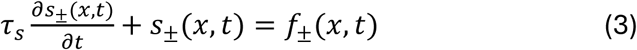

which describes the relaxation dynamics of the recurrent network, with a large timescale τ_*s*_= 80 ms compared to single neuron dynamics. The imbalance of inputs at non-zero velocity between the two rings makes one bump larger than the other (Figure 6C), and move both bumps with a speed proportional to *v* (see Figure 3d in Xie et al.^52^).

### Energy consumption rates

The energy rate at time *d* used by the CAN to generate action potentials is defined proportional to the sum of the neuronal firing rates in each ring: *E*^*AP*^(*t*) ∝ ∫ *dx* (*f*_−_(*x*, *d*) + *f*_+_(*x*, *d*)). Similarly, we consider the energy rate *E*^*ST*^(*t*) associated with synaptic transmission. We need to separate out in Eq.(2) the inhibitory and excitatory postsynaptic currents. The coupling matrices are decomposed into excitatory (positive) and inhibitory (negative) parts through *W* = *W*^*E*^ + *W^I^* with *W*^*E*^ = *W*_0_ + *W*_1_ and *W^I^* = *W*_1_(1 −*cos* (θ)). The full expression for the power due to synaptic transmission in the − ring is thus

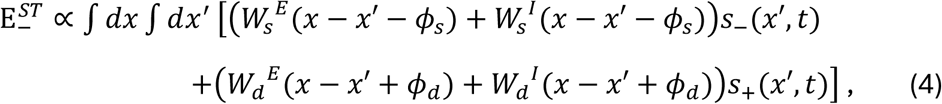

together with a similar expression for the rate 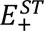 in the + ring. The sum of the two ST rates and the AP rates are shown in Figure 6D to increase quadratically with velocity.

We have verified the model-independence of our results by carrying out the same investigation on different theoretical models to good agreement. Notably, we carried out our analysis on a computational model of the Drosophila^95^, which is known to have both experimental and computational support. As seen in Supplementary Figure 10A,B, this model increases the peak firing rate, the height of the bump, and recruits more neurons to code for head direction, widening the bump, with increasing angular speed, which results in a supra-linear scaling between energy and speed. The scaling is not exactly quadratic owing to the fact that the head-direction circuit in the fly is modeled with 54 neurons only, and the recruitment of neurons saturates as velocity increases.

### Comparison with CBV data

Cerebral blood volume (CBV) and the cerebral metabolic rate of oxygen (CMRO_2_), a proxy for neuronal energy consumption, are both governed by cerebral blood flow (CBF). Griffeth and Buxton^96^ showed that CBV ∝ CBF^0.38^, and Buxton and Frank^97^ showed that CMRO_2_ ∝ CBF^0.84^. By combining these scaling laws, we assumed that Energy ≈ CMRO_2_ ∝ CBV^2.2^. This relationship varies slightly depending on the analysis and methods in question, but generally the exponent can range from 2 to 2.5. We found that the effect of this exponent range on the data analysis is minimal, and we chose a squared relationship in our analysis for simplicity.

We carried out PCA on the approximately 1000 voxels and selected the five components so that more than 95 % of the variance is explained. We calculate a measure for the energy by taking the sum of the squared five components 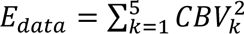. Finally, we normalize this by subtracting by the smallest value and then dividing by the standard deviation. We then define the sliding-averaged residual

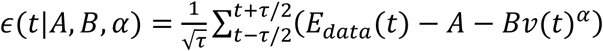

and the sum of squared residuals, *Q*(*A*, *B*, α) = ∑_*t*_ ε(*t*|*A*, *B*, α)^2^, over a session. We determine the optimal *A*, *B*, α that minimize *Q*.

In order to determine the appropriate sliding window size, we carried out a cross validation procedure, fitting the data to 80 % of the data and testing the residuals on the remaining 20 %. Performing this procedure, we found that the model starts to over fit when the window size is greater than 40 s. As can be seen in Supplementary Figure 9A, the size of the sum of the square residuals, Q, increases with increasing window size. However, for window sizes smaller than 10 s, the error in α increases greatly. In this plot, we define an effective error on α as the value of α at 0.5 % above the minimal Q. In other words, for small window size, the optimisation procedure finds an α with a shallow minimum that is less robust. Additionally, this is compatible with the predicted haemodynamic response trace which had a dynamics lasting approximately 10 s.

In order to verify that we are not fitting our model to noise, we carried out this optimization procedure for different permutations of the CBV data in time. Carrying out 500 randomized trials, we find that the model is not able to fit anything reasonable apart from the mean of the data. As can be seen in Supplementary Figure 9B, the optimal α is found to be around 0, the optimal A is found to be 1 and the optimal B is found to be zero, meaning that there is no correlation between velocity and CBV^2^ for shuffled data. Additionally, we performed the optimization procedure on white matter and caudate putamen, brain regions which we did not see any correlation with animal speed during the lagged analysis. As can be seen in Supplementary Figure 9C almost all fits showed either α or B close to 0.

### Animal tracking and behavior

The fUS imaging experiments were simultaneously recorded at 50 fps to allow for post-hoc pose estimation and animal tracking. For body part tracking we used DeepLabCut (version 2.3.2)^49,51^. We labeled 500 images taken from 13 different videos and 5 different rats. An EfficientNet-based^98^ neural network was trained for 50k iterations in 90% of the labeled frames. Train and test error was 1.59 pixels and 1.95 pixels respectively. The setup was designed to output synchronized CBV time series for each voxel in the selected imaging plane and any behavioral signals that can be extracted from the pre-trained labels in the pose estimation model^49–51^ (Figure 1A-C, Supplementary Figure 4). Grid-like and place-like expected activity were also computed by generating a large set of 2D functions representing theoretical grid and place fields and using the *x, y* instantaneous position of the animal as inputs (Supplementary Figure 5).

## Author contributions

SP, MT and FCP designed the experimental paradigm. NYR performed the surgical procedures and the animal’s care. FCP performed the ultrasound experiments and processed the ultrasound data. MT and BFO supervised the signal processing of the ultrasound data. SHC, SC and RM proposed the analysis of the computational model. FCP, SMD and SHC developed the code base needed for signal processing and data analysis. FCP, SHC, SC, RM, SP, MT were involved in the interpretation of the data and wrote the manuscript. All authors read and corrected the final version of the manuscript.

## Acknowledgments

This work was supported by Iconeus (FCP PhD fellowship), the ANRT (FCB PhD fellowship), the Institut National de la Santé et de la Recherche Médicale and the European Research Council (ERC) Advanced Grant FUSIMAGINE and the ESPCI.

Authors would like to thank Mrs L. Antoinette for animal husbandry. Finally, this work was supported by the Inserm ART (Technology Research Accelerator) in “Biomedical Ultrasound”.

Several panels of figures were made using Biorender (Biorender.com).

## Data availability statement

Source data will be available on a Zenodo repository (DOI: 10.5281/zenodo.15476373) which is currently private.

## Code availability statement

Custom codes used for the analysis of fUS data used in this study are protected by INSERM. Code used for the behavioral analysis will be available on a currently private Zenodo repository (10.5281/zenodo.15494995). The rest of this work did not involve any other particular code.

## SUPPLEMENTARY FIGURES

**Supplementary Figure 1:**
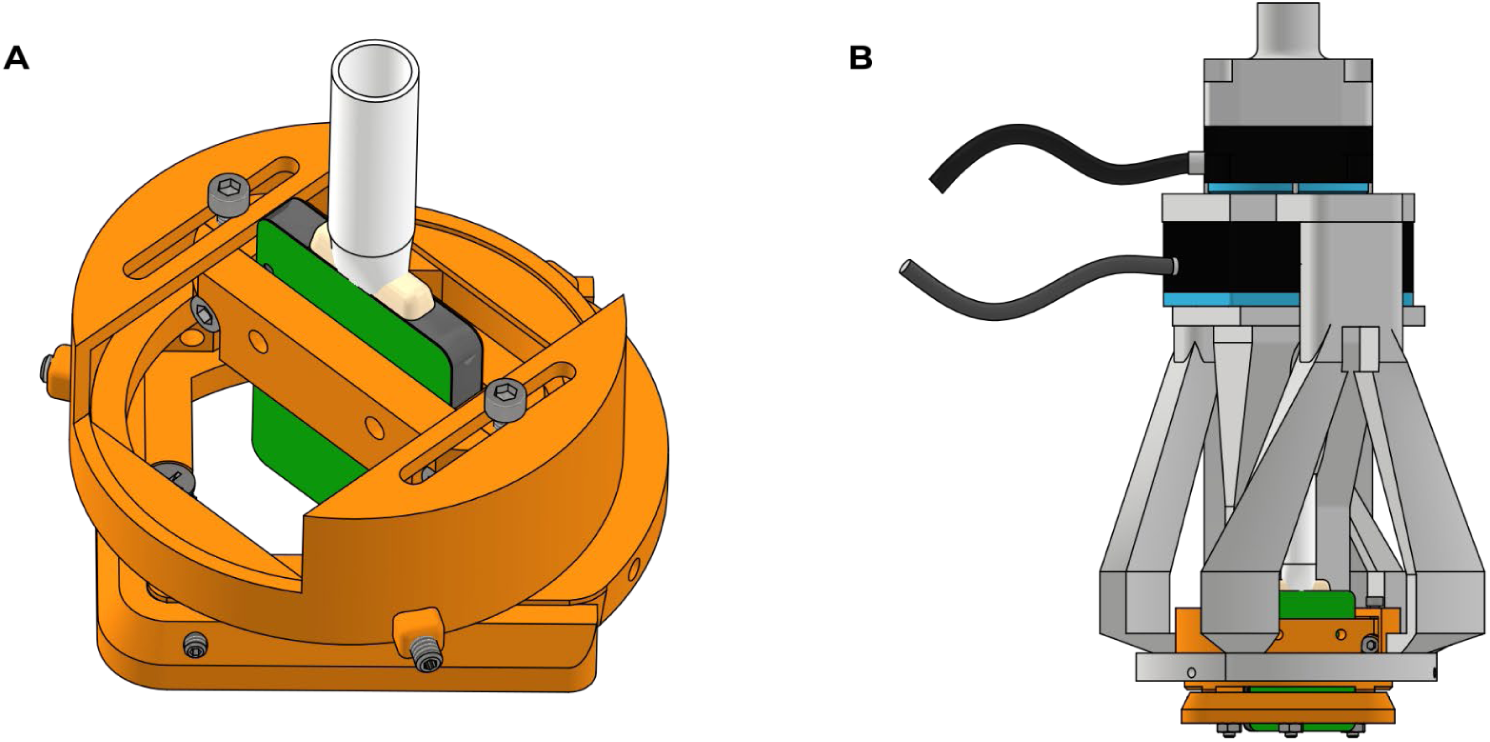
Schematic drawing of 3D printed ultrasonic probe holder and the motorized setup allowing neuronavigation with freely moving probe holder: A. 3D printed helmet that is screwed over the rat’s skull. The ultrasonic probe holder slides on top of the helmet and holds the probe via 2 pressure pieces. The probe holder has a rotation and a linear translation degree of freedom before fixation. **B.** 2-axis motorized setup that allows controlling the 2 degrees of freedom in the probe holder during anesthetized sessions.

**Supplementary Figure 2:**
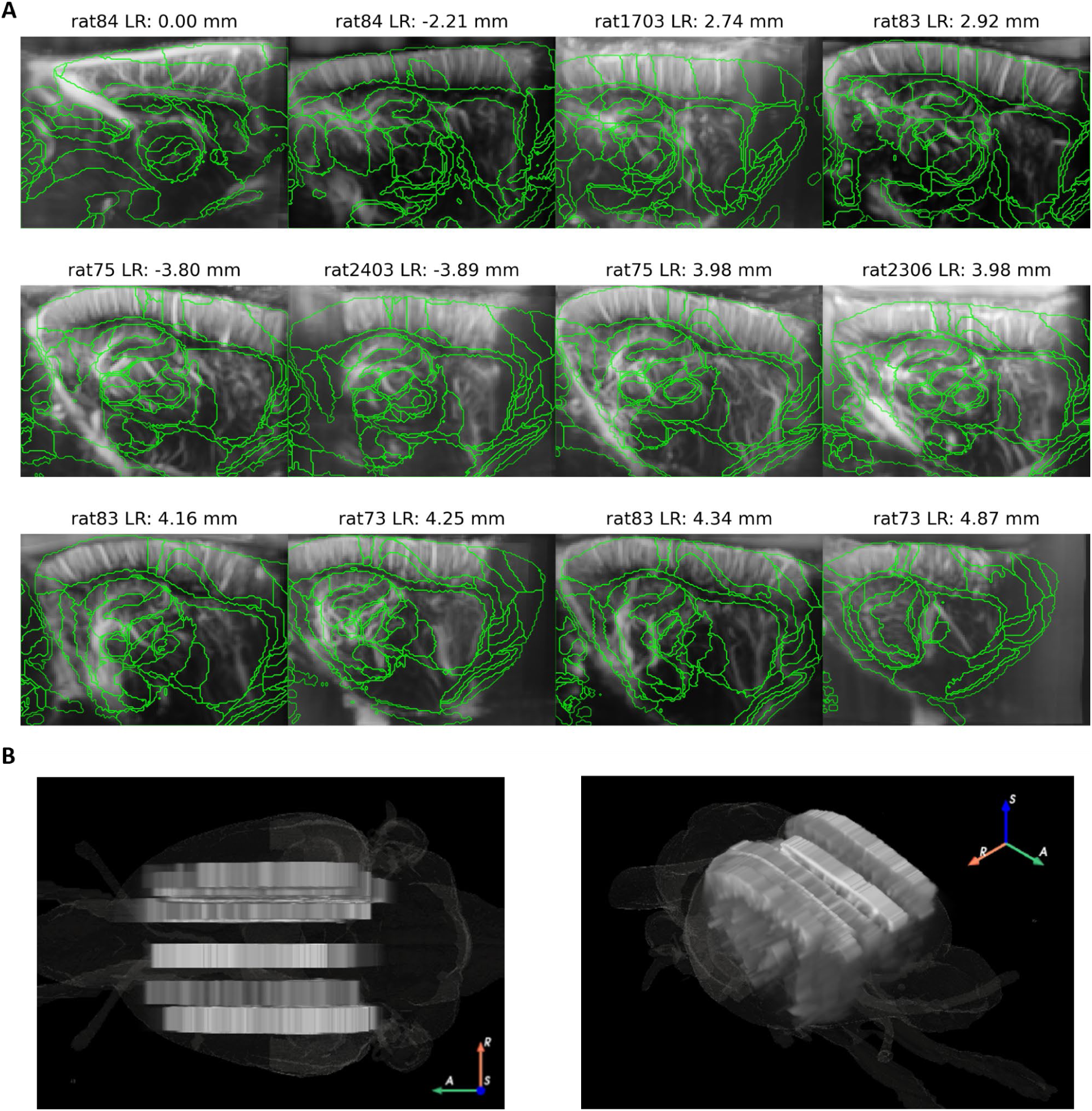
Imaging planes and anatomical atlas overlay across animals: A. Parasagittal imaging planes for all animals with registered anatomical atlas overlaid in green. Imaging planes are ordered from less to most distant of the sagittal midline. Titles show the distance from midline in a Left-Right (LR) scale. Midline is 0.00 mm, parasagittal to the left is negative, parasagittal to the right is positive. **B.** Volumetric view of all imaging planes in a RAS+ orientation. Left panel: yx view. Right panel: isometric view.

**Supplementary Figure 3:**
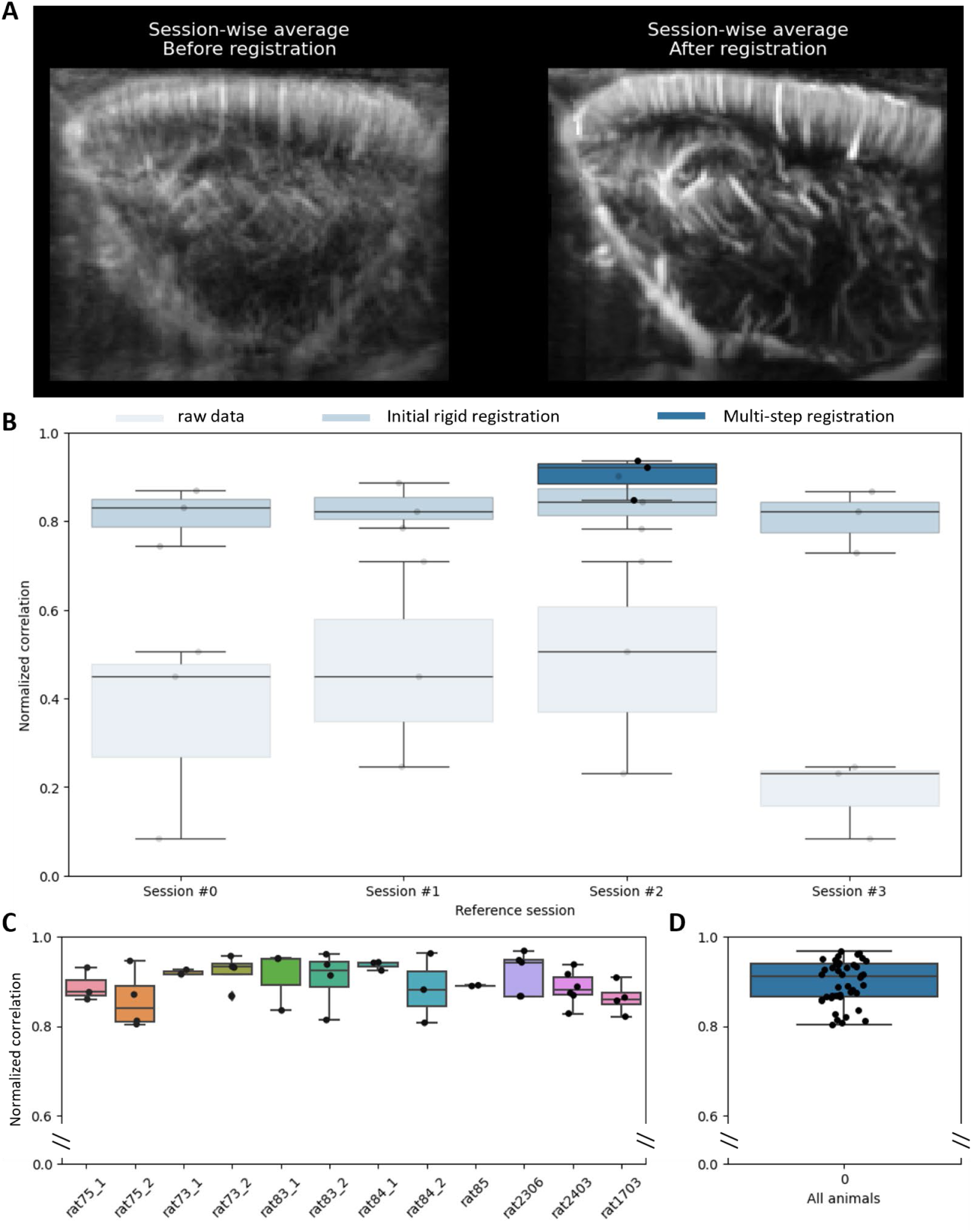
Intra-animal (inter-session) registration metrics for before-after registration comparison: A. Session-wise average before and after registration for the 4 sessions of an example rat. **B.** Normalized correlations between one of the sessions (reference session) and the rest of the sessions. The correlations using the raw data (no registration) are depicted in very light blue. The correlations using an initial rigid registration are depicted in light blue. The correlations using the multi-step registration are depicted in blue. The multi-step registration is only performed using the reference session that presented the highest correlation score in the initial rigid registration. **C.** The correlation metric for the multi-step registration method across all animals and imaging planes. **D.** All animals and imaging planes plotted in **C** in the same box-and-whisker plot. Box-and-whisker plot have whiskers defining the median, IQR and 1.5*IQR. Each dot represents one imaging session.

**Supplementary Figure 4:**
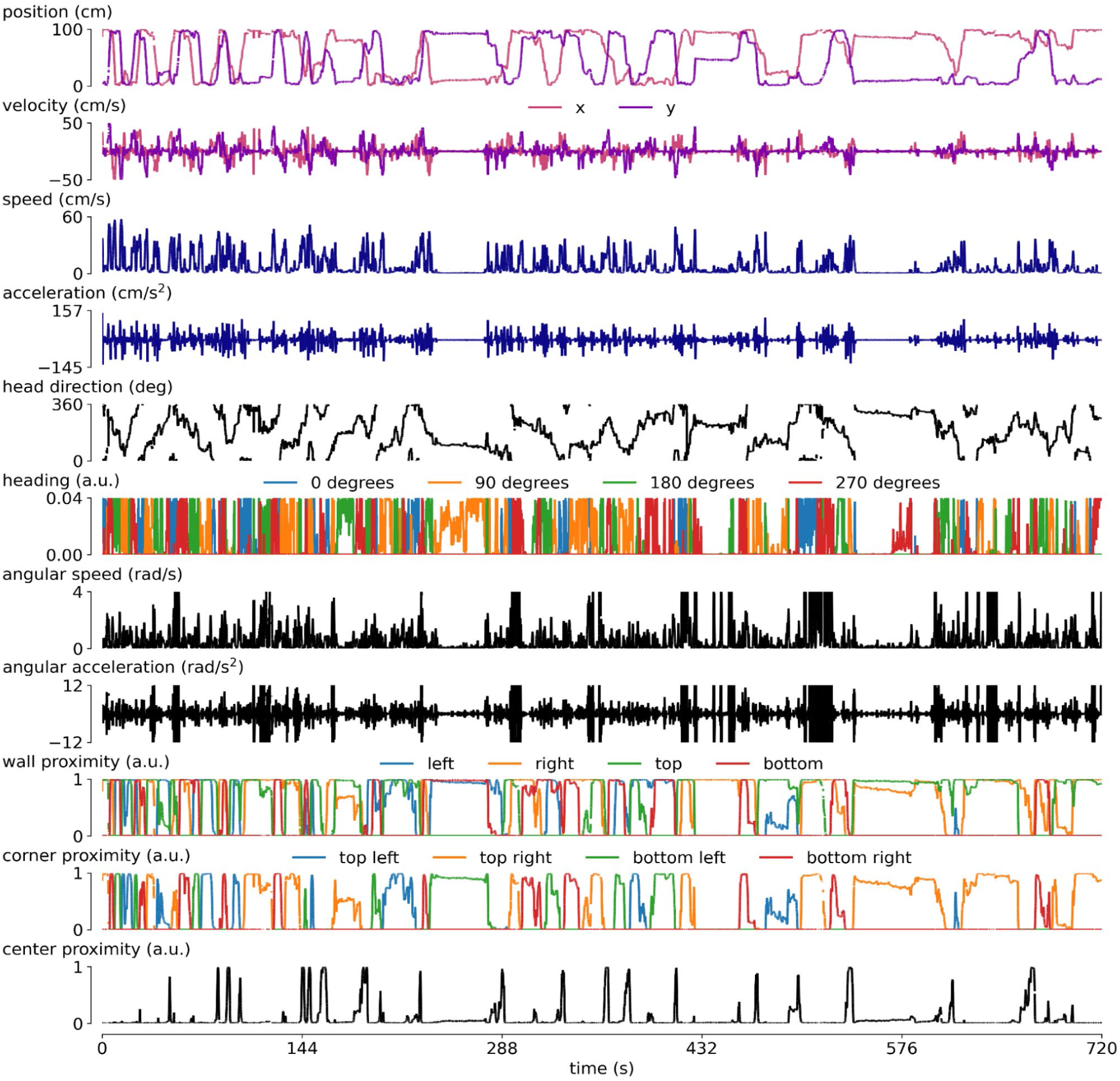
Different behavioral parameters extracted from DeepLabCut pose estimation: Plottings for different behavioral parameters extracted from the pose estimation using 5 labels in DeepLabCut (nose, probe, neck, body, tail). Positional parameters: position in the arena, wall proximity, corner proximity, center proximity. Speed parameters: velocity, speed, acceleration, angular speed, angular acceleration. Heading parameters: head direction, specific headings.

**Supplementary Figure 5:**
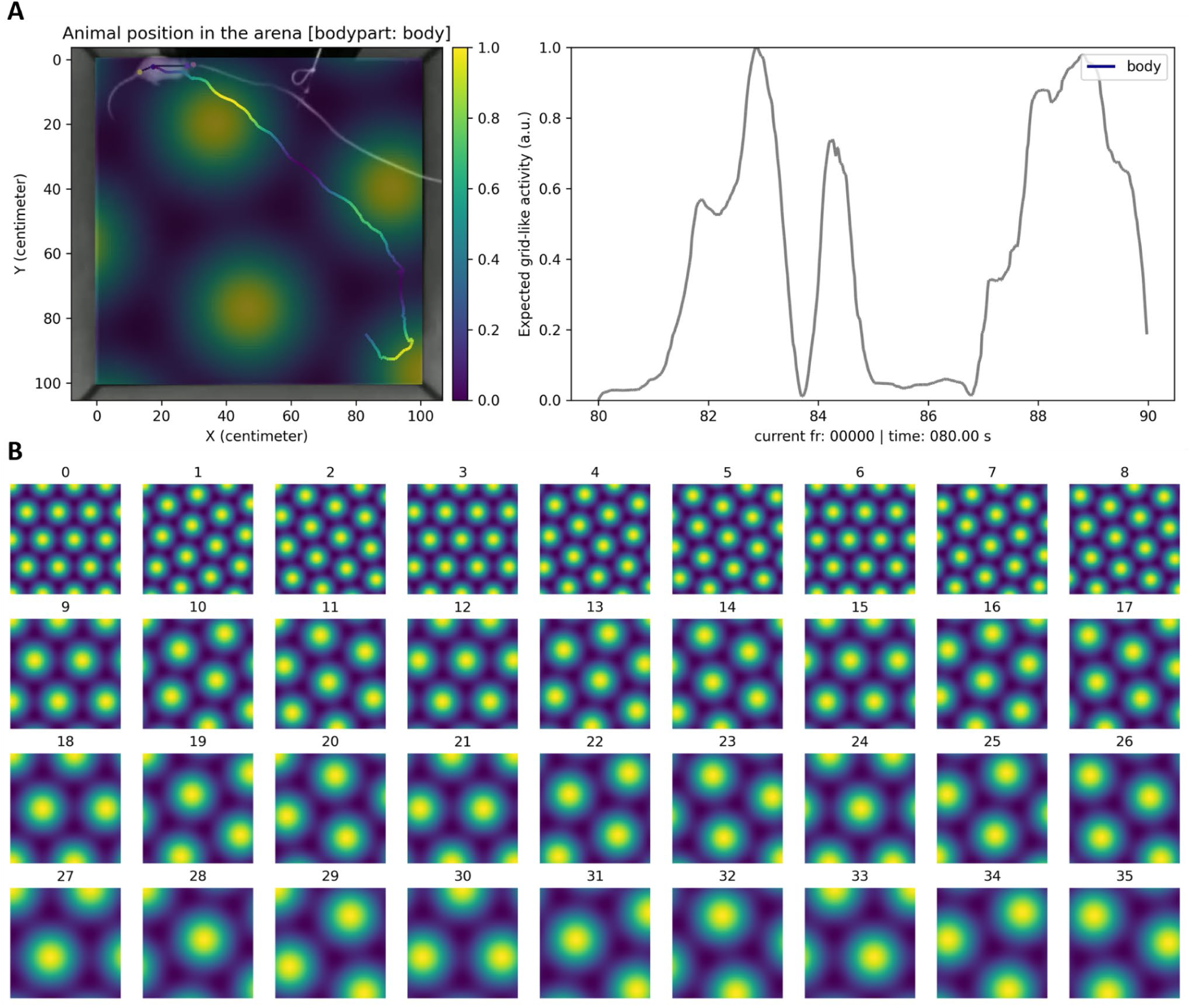
Simulation and computation of expected grid-like activity: A. Example representation of an expected grid-like activity for a given set of grid parameters. Left: Video frame of the animal in the arena overlaid by the theoretical grid field and the tracking position of the animal colored by the expected grid-like activity in this given grid field. Right: Plot of the expected grid-like activity during the time period depicted on the video frame. **B.** Full range of grid fields generated by the set of 36 different combinations of grid parameters used in this study.

**Supplementary Figure 6:**
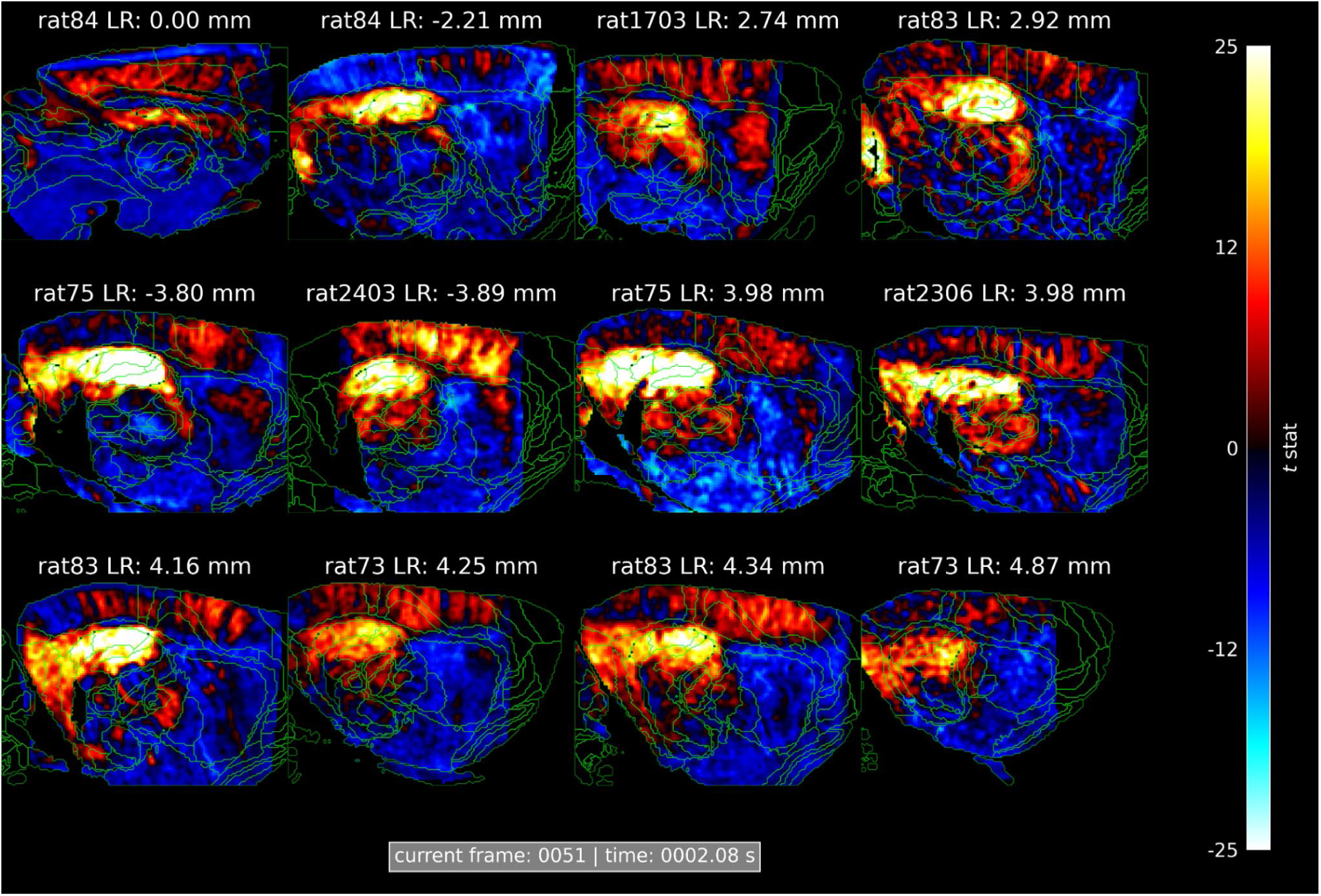
Peak activation for all imaging planes in the lagged GLM analysis for animal speed: Lag for which the activation in the hippocampal region is at its maximum across all animals and imaging planes for the lagged GLM analysis for animal speed.

**Supplementary Figure 7:**
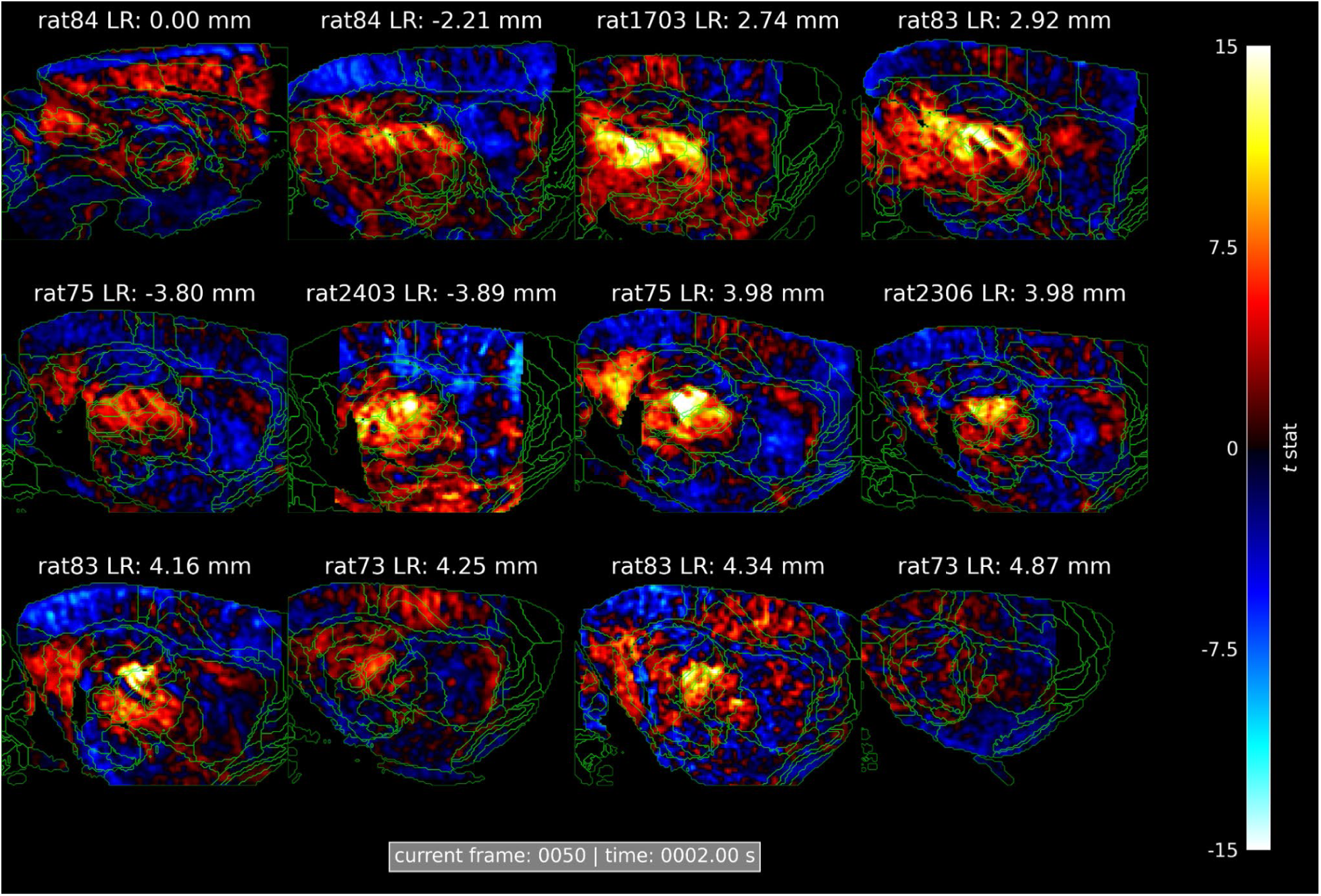
Peak activation for all imaging planes in the lagged GLM analysis for angular head speed: Lag for which the activation in the thalamic region is at its maximum across all animals and imaging planes for the lagged GLM analysis for head angular speed.

**Supplementary Figure 8:**
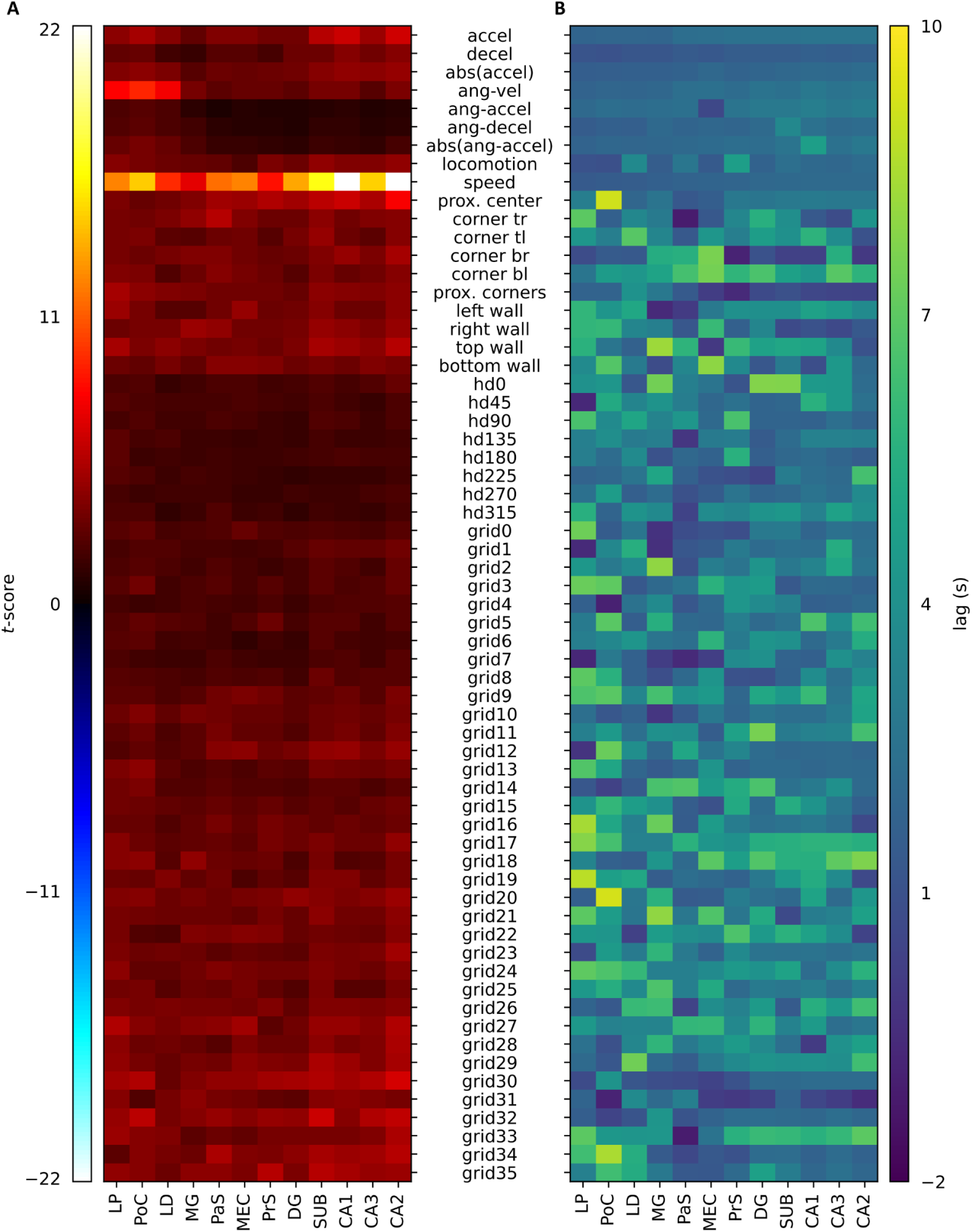
Summary lagged GLM results for all signals of interest tested: A. Median t-score value across animals for given ROIs in Figure 2 (x-axis) for each signal of interest tested with the lagged GLM approach. “accel” means positive acceleration, “decel” means deceleration or negative acceleration, “abs(accel)” means absolute acceleration value, “ang-vel” means angular velocity, “ang-accel” means angular positive acceleration, “ang-decel” means angular deceleration, “abs(ang-accel)” means absolute angular acceleration value, “locomotion” means binary moving-not moving signal, “speed” means animal locomotion speed, “prox. center” means proximity to the center of the open field, “corner tr” means proximity to the top right corner, “tl”, “br”, and “bl” means top left, bottom right and bottom left, respectively, “prox. corners” means proximity to the corners of the open field, “left, right, top and bottom wall” means proximity to the corresponding wall, “hd{i}” means head direction at i degrees, “grid{i}” means gridlike pattern number i (see Supplementary Figure 5) **B.** Median lag to peak for the same ROIs and signals of interest.

**Supplementary Figure 9:**
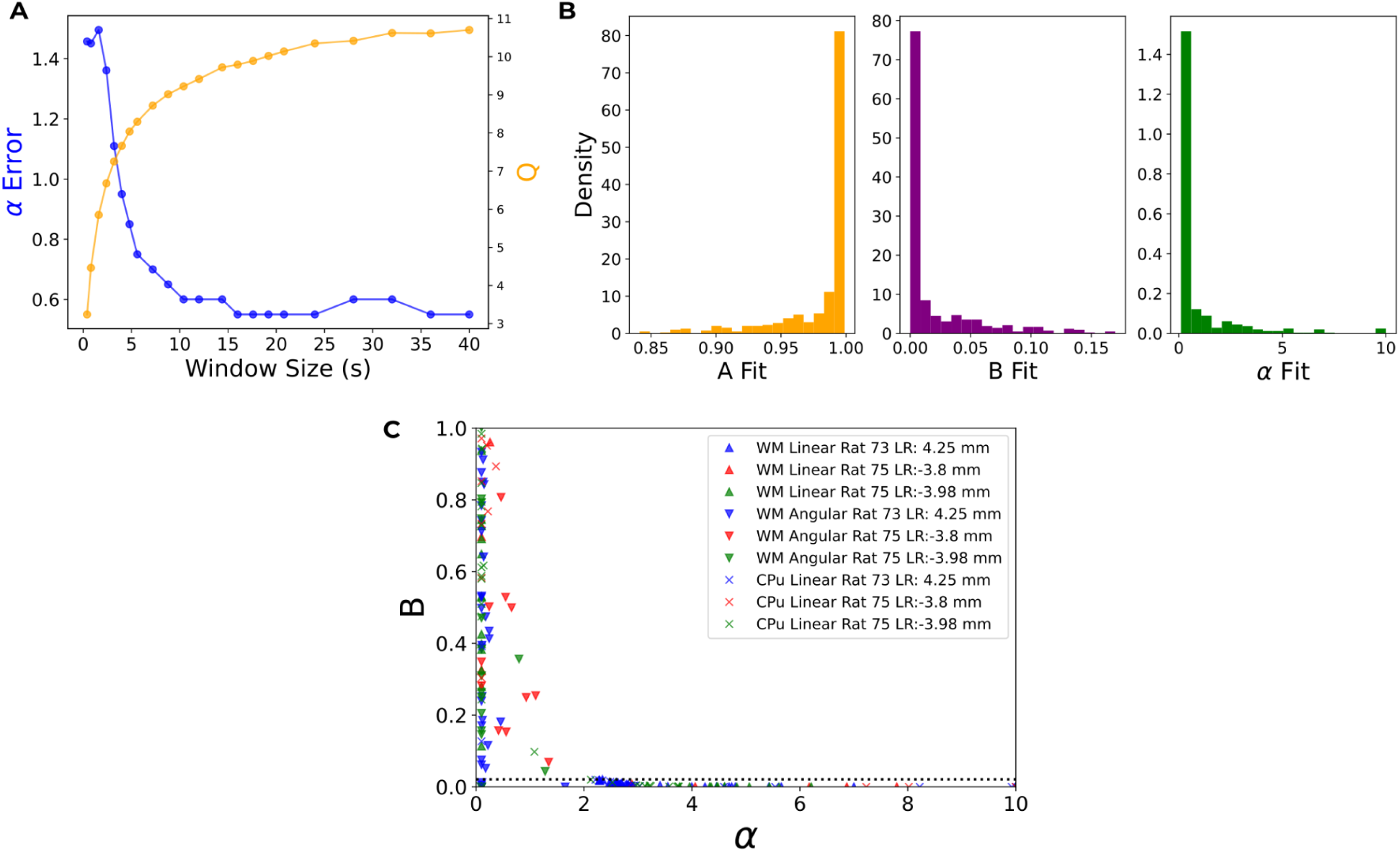
Cross-Validation and null tests confirm model robustness and unique hippocampal/thalamic signals, avoiding noise overfitting: A. Dependence of the error on α and the residual sum of squares, Q, on the sliding average window size. An optimal window of 10 s minimizes Q while keeping the uncertainty on α low. **B.** Distributions of best-fit parameters obtained from 500 random permutations of the CBV^2^ data. The parameter values show no correlation between energy and velocity. This confirms that the fitting procedure does not overfit noise. **C.** This panel shows the result after optimisation of data that we expect not to have any correlation. This includes correlation between white matter (WM) and linear or angular velocity and correlations between the caudate putamen (CPu) and linear velocity. Nearly all optimisations result in either small B or small α. This shows that the correlation we see in the hippocampus and thalamus are due to signals that are unique to that brain area.

**Supplementary Figure 10:**
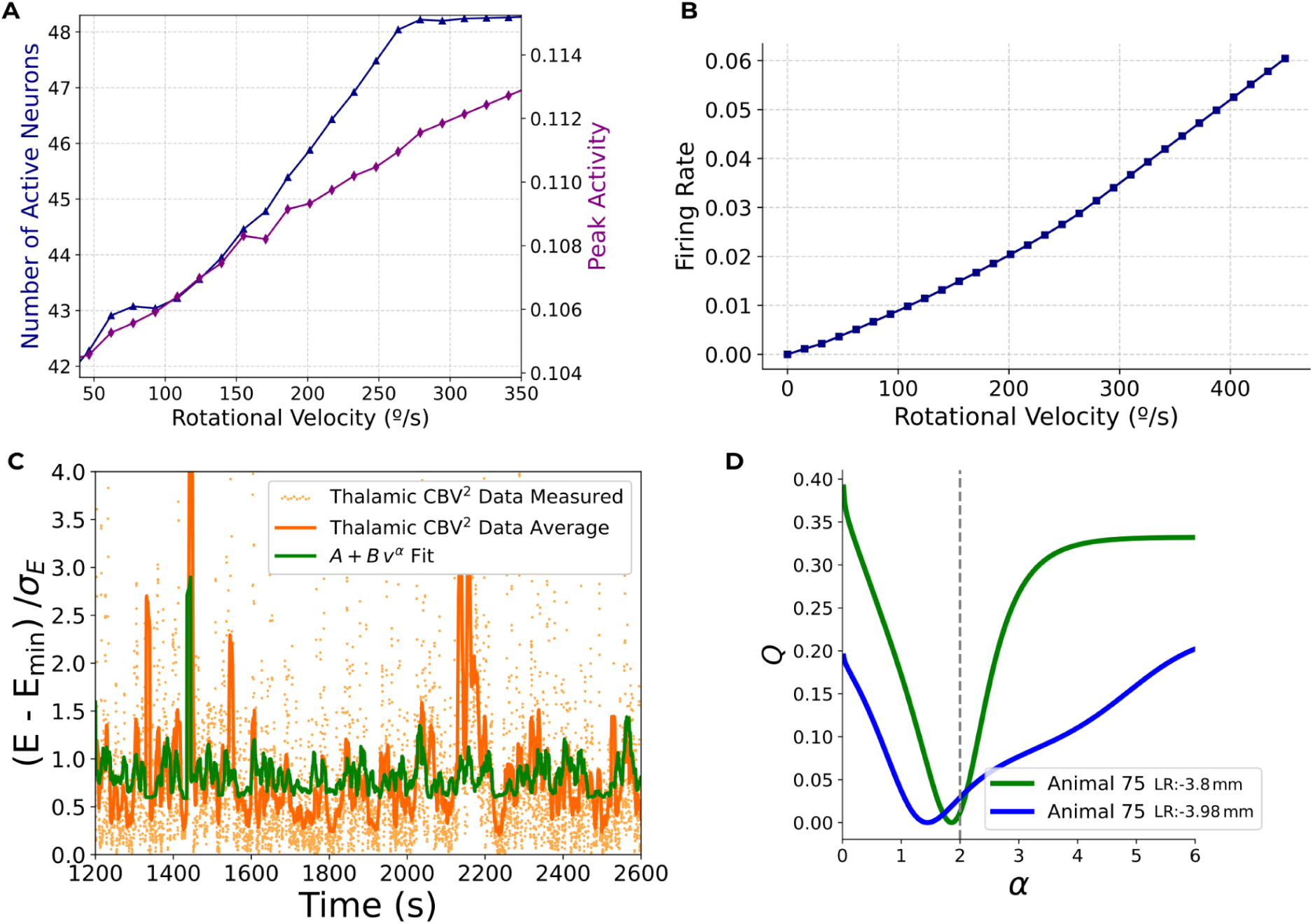
Head direction modelling -Conceptual agreement with a computational model of Drosophila and experimental validation of head direction signal in CBV data: A. This panel shows the dependence of the number of active neurons and peak activity on angular velocity in a computational model of the Drosophila head-direction system^95^. Both quantities increase linearly with velocity, resulting in a supra-linear relationship between firing rate and velocity, as illustrated in panel **B**. A power-law fit yields an exponent of 1.3. This sub-quadratic scaling, compared to that of the larger model given above, reflects the saturation effects due to the small number of neurons (∼50). **C.** CBV2 time series in the thalamus for animal 75 (LR:-3.8 mm). Orange points indicate the raw measurements; the orange line is a 10 s moving average. The green line shows the corresponding best fit of the model *A* + *Bv*^α^, where *v* is the animal’s angular velocity. **D.** This panel displays the residual Q (after optimization over A and B) as a function of α for selected sessions from two slices.

